# Growth Signaling Autonomy in Circulating Tumor Cells Aids Metastatic Seeding

**DOI:** 10.1101/2022.12.02.518910

**Authors:** Saptarshi Sinha, Alex Farfel, Kathryn E. Luker, Barbara A. Parker, Kay Yeung, Gary D. Luker, Pradipta Ghosh

**Affiliations:** Department of Cellular and Molecular Medicine, School of Medicine, University of California San Diego, La Jolla, CA 92093; Biointerfaces Institute, University of Michigan.; Moores Cancer Center, University of California San Diego, La Jolla, CA 92093; Department of Medicine, School of Medicine, University of California San Diego, La Jolla, CA 92093; Department of Biomedical Engineering, University of Michigan, 109 Zina Pitcher Place, Ann Arbor, MI, 48109-2200, USA; Department of Microbiology and Immunology, University of Michigan, 109 Zina Pitcher Place, Ann Arbor, MI, 48109-2200, USA; Center for Molecular Imaging, Department of Radiology, University of Michigan, 109 Zina Pitcher Place, Ann Arbor, MI, 48109-2200, USA; Veterans Affairs Medical Center, La Jolla, CA

**Keywords:** Cellular Autonomy, GIV, Girdin, CCDC88A, metastasis, mesenchymal-epithelial transition (MET), Epithelial-mesenchymal transition (EMT), Epithelial mesenchymal plasticity

## Abstract

Self-sufficiency (autonomy) in growth signaling, the earliest recognized hallmark of cancer, is fueled by the tumor cell’s ability to ‘secrete-and-sense’ growth factors; this translates into cell survival and proliferation that is self-sustained by auto-/paracrine secretion. A Golgi-localized circuitry comprised of two GTPase switches has recently been implicated in the orchestration of growth signaling autonomy. Using breast cancer cells that are either endowed or impaired (by gene editing) in their ability to assemble the circuitry for growth signaling autonomy, here we define the transcriptome, proteome, and phenome of such autonomous state, and unravel its role during cancer progression. We show that autonomy is associated with enhanced molecular programs for stemness, proliferation, and epithelial-mesenchymal plasticity (EMP). Autonomy is both necessary and sufficient for anchorage-independent growth factor-restricted proliferation and resistance to anti-cancer drugs and is required for metastatic progression. Transcriptomic and proteomic studies show that autonomy is associated, with a surprising degree of specificity, to self-sustained EGFR/ErbB signaling. Derivation of a gene expression signature for autonomy revealed that growth signaling autonomy is uniquely induced in circulating tumor cells (CTCs), the harshest phase in the life of tumor cells when it is deprived of biologically available EGF. We also show that autonomy in CTCs tracks therapeutic response and prognosticates outcome. These data support a role for growth signaling autonomy in multiple processes essential for the blood-borne dissemination of human breast cancer.

**GRAPHIC ABSTRACT:** 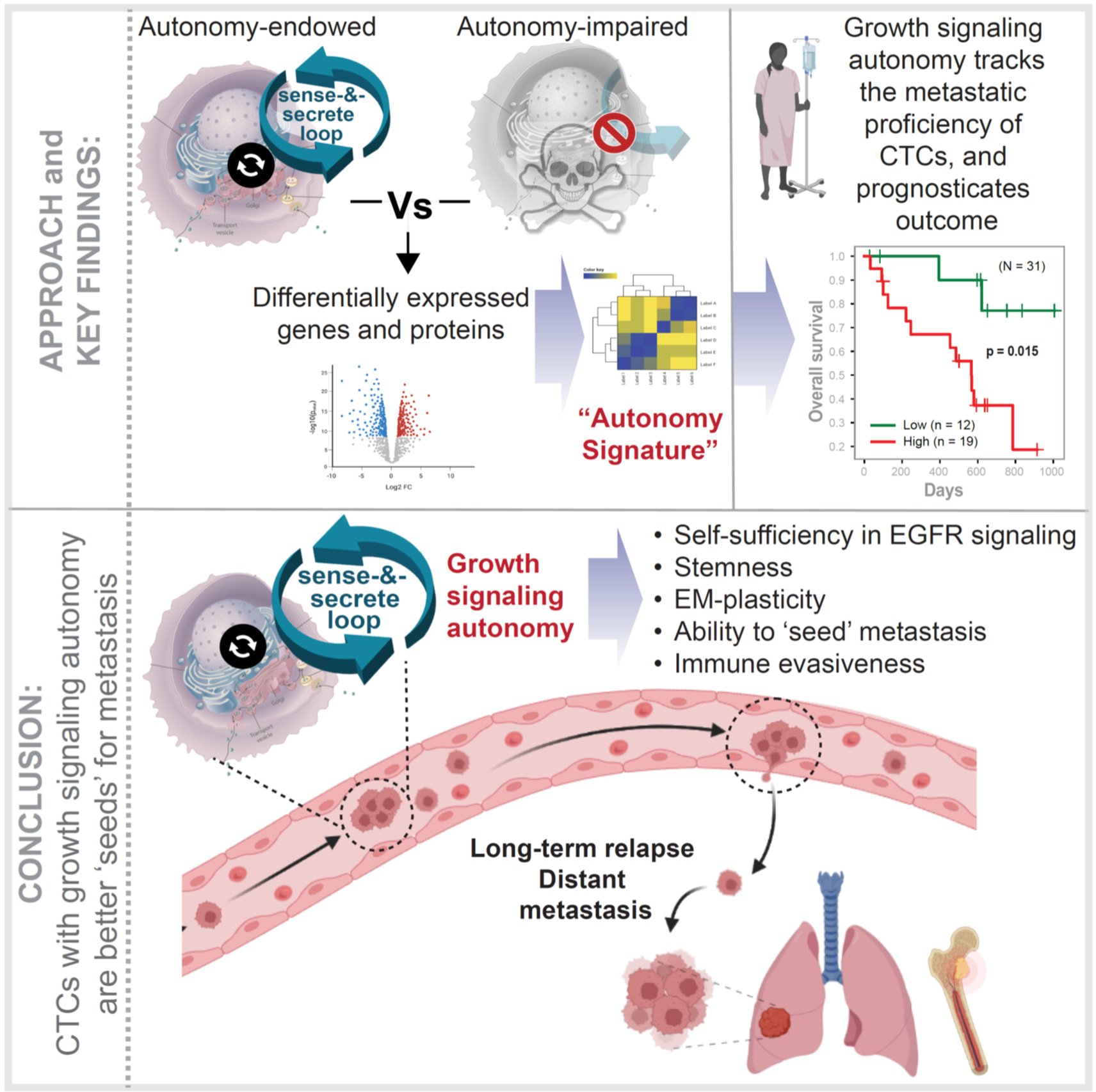

**Significance Statement:** A Golgi-localized molecular circuitry has been recently implicated in the orchestration of secrete-and-sense auto-/paracrine loops that impart self-sufficiency in growth signaling, a.k.a., growth signaling autonomy. Using a transdisciplinary approach, this work shows that growth signaling autonomy is uniquely induced in tumor cells that are in circulation. Circulating tumor cells (CTCs) represent a brutish and risky phase in the lifetime of tumor cells when they are exposed to the immune system and hemodynamic sheer forces, all in the setting of growth factor starvation. Cancer cells appear to rely on the autonomy circuit to survive and enhance their fitness to seed metastases. Autonomy generates the kind of ‘eat-what-you-kill’ entrepreneurial spirit which minimizes the risk of CTCs dying on an otherwise risky journey.

## Introduction

Metastatic breast cancer (MBC) remains a fatal disease. While ∼5-10% of patients are diagnosed with MBC at initial diagnosis, ∼20-30% of patients with stage I-III breast cancer will eventually recur with MBC. Hence, understanding which cells ‘seed’ metastases in MBC is of paramount importance to improve the management and outcome of these patients.

Metastasis begins with the intravasation of cancer cells from primary tumors, either as single cells or in clusters, into the systemic circulation. These circulating tumor cells (CTCs) must then extravasate from the blood stream and disseminate to distant tissues, where they either remain dormant or give rise to metastases (1–3). However, for CTCs to ‘seed’ metastases, they must survive despite the loss of anchorage to the matrix, exposure to the immune system and hemodynamic shear forces. The past two decades has witnessed significant technological leaps to help detect, enumerate, and characterize CTCs [reviewed in (4, 5)]. Numerous correlations have been discovered between CTCs and the ability to initiate metastases: abundance (6), phenotypic properties, e.g., enhanced protein translation (7), proliferation (7, 8), epithelial-mesenchymal plasticity (9–11), epithelial-type CTCs [with restricted mesenchymal transition (7, 12)] that assemble junctions and circulate as clusters (13–16), CTC ½ life (17), the presence of platelets (15) and immune cells [such as neutrophils (8)] in those clusters, hypoxia (18) and circadian rhythm [nighttime worse than day (19)]. Despite these insights, mechanisms connecting these diverse phenotypes to drive aggressive features of CTCs remain unknown. While clustering, either homotypic, with each other, or heterotypic with platelets provide explanation for how CTCs protect themselves from shear forces and evade the immune system, little to nothing is known about how they survive the journey in the face of a precipitous drop in growth factors. For example, the concentration of EGF within primary tumors and xenografts range from ∼1–2 ng/ml (20); however, serum EGF levels range between ∼13-14 pg/ml (21). Of the detectable EGF in serum, most of it is biologically inactive and is associated with platelets (22), which is released during the process of coagulation (23). In fact, EGF is virtually undetectable in plasma that is collected in the presence of inhibitors of coagulation (24).

It is perhaps because of these adversities that the process by which CTCs initiate metastases is highly inefficient [2.7% efficiency, as determined using MBC-patient derived CTCs in xenotransplantation models (25)], implying that only a fraction of CTCs--the fittest of them all--are endowed with tumorigenic and metastatic functionality. What signaling mechanisms and/or molecular machineries impart or maintain CTC fitness in growth factor-deprived state remains a hot topic of debate, and objective molecular measurements of the metastatic potential of CTCs remains an unattainable Holy Grail of precision medicine.

Here we report the serendipitous discovery of a distinct CTC phenotype, i.e., growth signaling autonomy that is induced in CTCs, but not in primary tumors or established metastases. Growth signaling autonomy, or self-sufficiency in growth factor (GF) signaling, is the first of the six hallmarks of all cancers to be defined (26), and yet remains one of the least well understood. Many cancer cells synthesize growth factors (GFs) to which they are responsive, creating a positive feedback signaling loop called autocrine stimulation (27). In fact, serum-free cell culture studies squarely implicate autocrine secretion of growth factors as key support for intracellular mechanisms that impart autonomy [reviewed in (28)]. Recently, using an integrated systems and experimental approach, a molecular circuit has been described which is critical for multiscale feedback control to achieve secretion-coupled autonomy in eukaryotic cells (29). This circuit is comprised of two species of GTPases, monomeric Arf1 and the heterotrimeric Gi, coupled by the multi-modular scaffold GIV, (i.e., *<u>G</u>*α-*i*nteracting *<u>v</u>*esicle associated protein; aka Girdin; gene *CCDC88A*) within a closed-loop circuit that is localized at the Golgi (29, 30) Coupling is initiated only when cells are subjected to restricted growth factor conditions. Coupling within such a closed-loop control system generates two emergent properties: (i) dose-response alignment behavior of sensing and secreting growth factors; and (ii) multiscale feedback control to achieve secretion-coupled growth and survival signaling (31). Consequently, cells with a coupled circuit are self-sufficient in growth signaling, i.e., autonomous, and can survive and achieve homeostasis in GF-restricted conditions; cells in which the circuit is uncoupled (as in GIV-KO cells) are not.

In this work, we provide evidence for the requirement of such autonomy in breast cancer CTCs and reveal the biological implications and translational potential of our observations.

## Results

### Study design

To study how autonomy in cancer cells impact cancer progression, and more specifically, breast cancers, we took the advantage of two MDA MB-231 breast cancer cell lines, that are either endowed (WT) or impaired (GIV-KO by CRISPR (29)) in growth signaling autonomy (**Fig 1a**). We focused on these cells because they are a highly aggressive, invasive, and poorly differentiated triple-negative breast cancer (TNBC) cell line which lacks the estrogen receptor (ER), progesterone receptor (PR), as well as amplification of the HER2 (human epidermal growth factor receptor 2) receptor and is one of the triple-negative basal subtype cell line most widely used in metastatic breast cancer research (32) (40.2% of total PubMed citations). It is also a cell line that has been shown to require GIV for growth signaling autonomy (29). We analyzed these cells by functional and ‘omics’-based approaches to navigate the uncharted territory of cancer cell autonomy. Because the GTPase circuit for autonomy requires GIV’s modules/motifs that evolved only in the higher eukaryotes (29), and GIV is overexpressed in most cancers (33), we hypothesized that tumor cells may frequently assemble and utilize such an evolutionary advantage to achieve growth signaling autonomy at some stage during cancer progression. Using an integrated computational and experimental approach, we systematically analyzed these pair of cell lines for key hallmarks of cancer cells.

**Figure 1.**
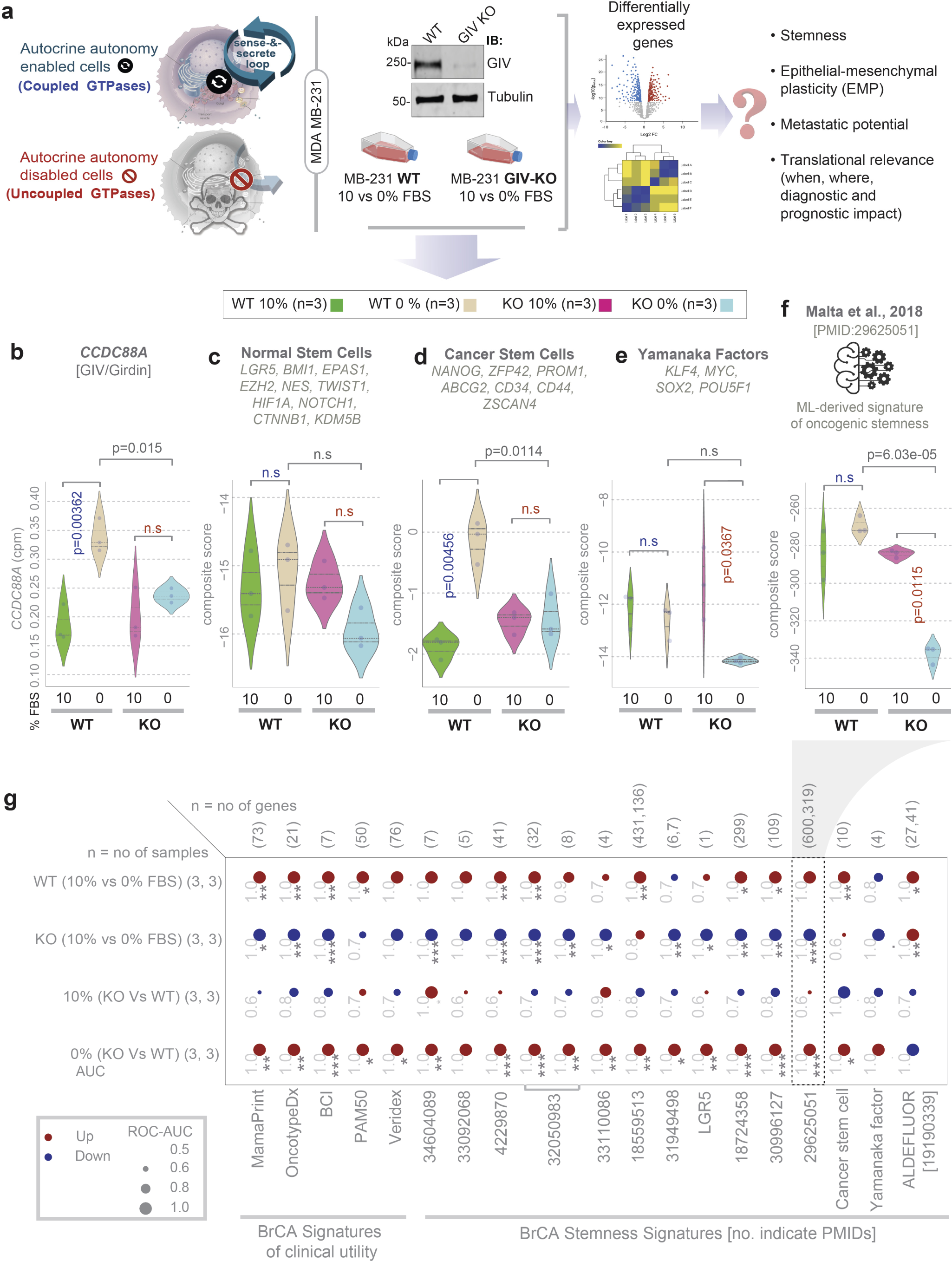
Growth signaling autonomy is required for the induction or maintenance of stemness. **a.** Schematic displays study design. Autocrine autonomy-endowed cells are compared against cells in which such autonomy is disabled (autonomy-impaired) by depletion of GIV (GIV-KO). Cells are grown in the presence or absence of exogenous growth factors (0% FBS) to study the biological and translational relevance of autocrine autonomy in breast cancers. **b-f.** Violin plots display the single (b) or composite (c-f) score of selected gene signatures. P values based on Welch’s T-test, comparing 10% vs 0% growth conditions in WT (blue) and KO (red) cells. Blue and red font for p values indicate, significant up- or downregulation, respectively. **g.** Expression of various stemness-associated gene signatures and clinically used breast cancer gene signatures, in parental (WT) vs GIV-KO (KO, by CRISPR) MDA MB231 cells grown in 10% or 0% FBS is visualized as bubble plots of ROC-AUC values (radius of circles are based on the ROC-AUC) demonstrating the direction of gene regulation (Up, red; Down, blue) for the classification of WT and KO samples in 10% and 0% FBS conditions based on the indicated gene signatures (bottom). BCI, breast cancer index. Nos. indicate PMIDs. *Statistics*: *p* values based on Welch’s T-test (of composite score of gene expression values) are provided either as exact values (in panels b-f) or using standard code in panel g (‘’p>0.1, ‘.’p<=0.1, *p<=0.05, **p<=0.01, **p<=0.001) next to the ROC-AUC. n.s., not significant. See **Figure S1** for violin plots of representative signatures.

### Growth signaling autonomy is associated with stemness and epithelial-mesenchymal plasticity

The expression of *CCDC88A* gene (which encodes GIV) was significantly upregulated when the autonomous WT, but not GIV-KO cells were switched from 10% to 0% serum conditions (**Fig 1b**), which is consistent with the increased need for autocrine growth factor signaling during serum starvation (absence of exogenous growth factors, i.e., 0% FBS). Although conventional markers of normal pluripotent stem cells remained unchanged during serum depravation in both cells (**Fig 1c**), markers of cancer stem cells (CSCs) followed the same pattern as *CCDC88A* (**Fig 1d**). When we analyzed the core pluripotency master regulators (the Yamanaka factors (34); **Fig 1e**) and the breast cancer-specific indices of the degree of oncogenic dedifferentiation (identified using a machine learning algorithm (35)) (**Fig 1f**), the autonomous WT cells maintained these signatures despite serum depravation; GIV-KO cells did not. These patterns (induction or maintenance in the autonomous WT, but suppression in GIV-KO cells) were observed repeatedly across a comprehensive panel of gene signatures of breast cancer aggressiveness and stemness that have been reported in the literature (**Fig 1g**; **Figure S1**).

Autonomous WT, but not GIV-KO cells also induced gene signatures for epithelial-mesenchymal plasticity (EMP (36)), i.e., the ability of cells to interconvert between epithelial and mesenchymal phenotypes in response to signals (**Fig 2**). For example, all gene signatures derived from isolated distinct single-cell clones from the SUM149PT human breast cell line spanning the E↔EM1-3↔M1-2 spectrum, previously characterized for diverse migratory, tumor-initiating, and metastatic qualities (37), were induced in the autonomous WT, but suppressed in the GIV-KO cells during serum depravation (**Fig 2a-b**). Acquisition of EMP plasticity in MDA-MB-231 cells under conditions of serum deprivation is remarkable since these cells typically show a highly mesenchymal phenotype(38). An identical pattern was seen also for the transcriptional census of EMP in human cancers, which was derived by leveraging single-cell RNA seq data from 266 tumors spanning 8 different cancer types (39) (**Fig 2c**). This held true for both the 328-gene EMP consensus signature (**Fig 2d**), as well as its cancer cell-specific 128-gene subset (**Fig 2e**). Furthermore, numerous gene signatures across the EMT and MET spectrum, derived from diverse human samples representing the stages of metastasis (primary tumors, CTCs, and metastases) and genes that are essential for establishing cell-cell junctions were induced in the autonomous WT, but remained unchanged or were suppressed in the GIV-KO cells (**Fig S2**).

**Figure 2.**
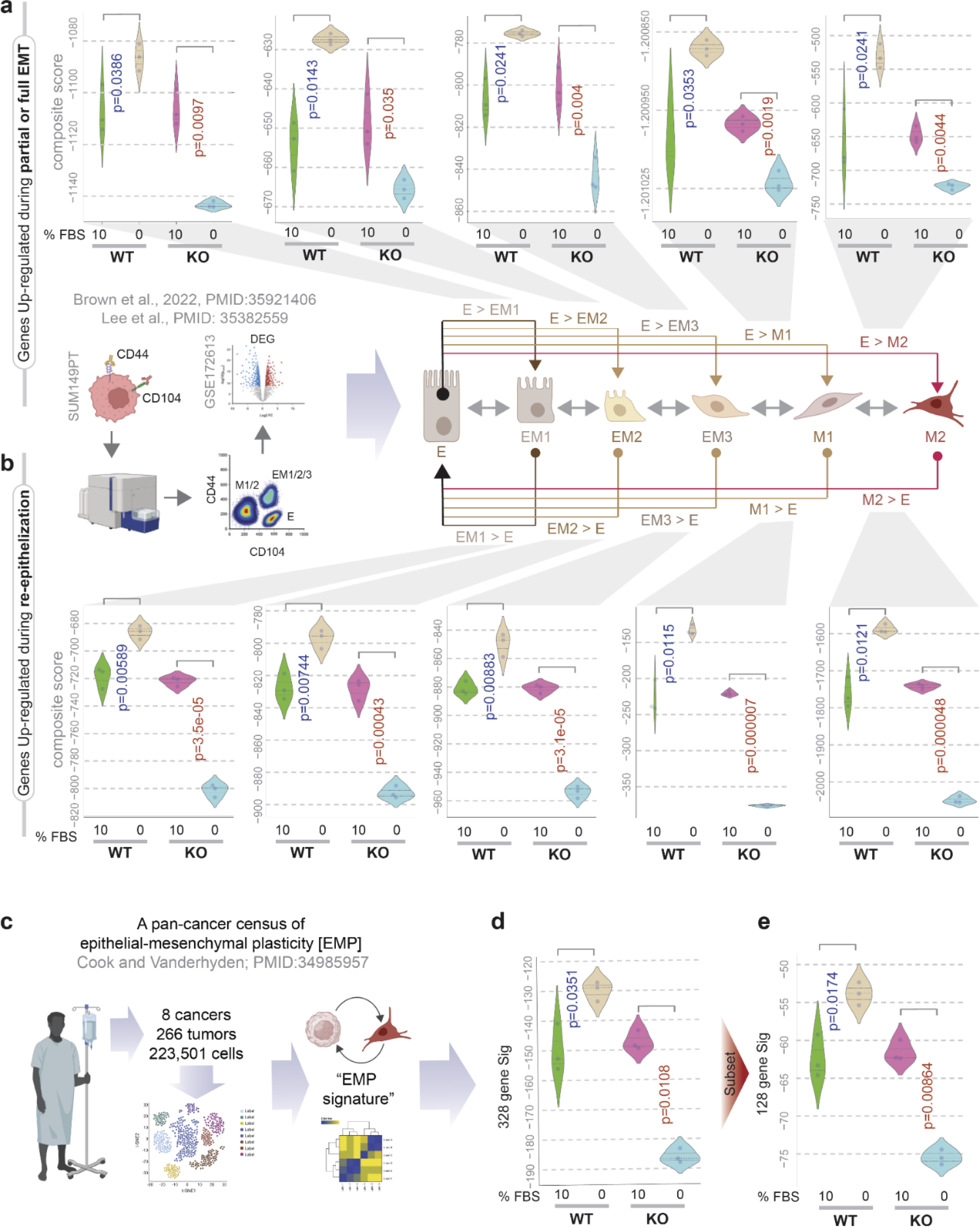
Growth signaling autonomy is required for the induction of molecular programs for EMT, re-epithelization and epithelial-mesenchymal plasticity (EMP) **a.** Violin plots display the composite score of various gene signatures in parental (WT) and GIV-depleted (GIV-KO) MDA MB-231 cells, which are induced during epithelial cell transition into partial (EM1, EM2, EM3) or full M1 and M2) EMT states. **b.** Violin plots display the composite score of various gene signatures in parental (WT) and GIV-depleted (GIV-KO) MDA MB-231 cells, which are induced during the re-epithelization of partial (EM1, EM2, EM3) or full (M1 and M2) EMT states. **c-e**. Schematic (c) summarizes the study design that resulted in the discovery of the pan-cancer transcriptional census of epithelial mesenchymal plasticity (EMP) derived by single cell seq (39). Violin plots (d-e) display the composite score of the EMP signature in parental (WT) and GIV-depleted (GIV-KO) MDA MB-231 cells. Panel d shows the expression profile of all 328 genes in the EMP signature, representing various cells in the tumor microenvironment. Panel e shows the expression profile of a subset of 128 genes in the EMP signature that is specifically enriched in cancer cells. *p* values based on Welch’s T-test, comparing 10% vs 0% growth conditions in WT (blue) and KO (red) cells. Blue and red font for p values indicate, significant up- or downregulation, respectively. See **Figure S2** for other gene signatures for EMT, CTC clusters and MET.

These findings indicate that GIV-dependent growth signaling autonomy is required to support molecular programs of stemness and EMP in growth factor-restricted conditions (i.e., 0% FBS); however, such autonomy is largely dispensable in the presence of excess growth factors because all readouts were indistinguishable when WT and KO cells were compared at 10% FBS (**Fig 1-2**).

### Autonomy is required for anchorage-independent growth and metastatic spread

We next assessed if the autonomy-endowed and- impaired cells are capable of anchorage-independent growth, which is a hallmark of anoikis resistance and the path to further steps in metastasis (40). When tested for growth as spheroids in soft agar at varying serum concentrations, while both WT and GIV-KO cells did so in the presence of excess serum, only the autonomous WT cells thrived in serum-restricted conditions (0.2% FBS; **Fig 3a-c**). Under serum-restricted growth conditions, the autonomous WT, but not the autonomy-impaired GIV-KO cells were also relatively resistant to various classes of conventional chemotherapeutic agents that are typically used to treat TNBCs (e.g., anthracyclines, alkylating agents), as determined by the observed differences in their half-maximal inhibitory concentrations (IC_50_; **Fig 3d**). Compared to the KO cells, the autonomous WT cells also displayed significantly higher metastatic potential after intracardiac injection (**Fig 3e-f**). Notably, WT and GIV-KO cells produced comparable metastases when cells were cultured in complete serum prior to intracardiac injection (compare WT and KO, 10% FBS; **Fig 3f**). These findings indicate that GIV is required for 3D growth, chemoresistance and metastasis in serum-restricted conditions and that GIV-dependent growth signaling autonomy may be required for these phenotypes.

**Figure 3.**
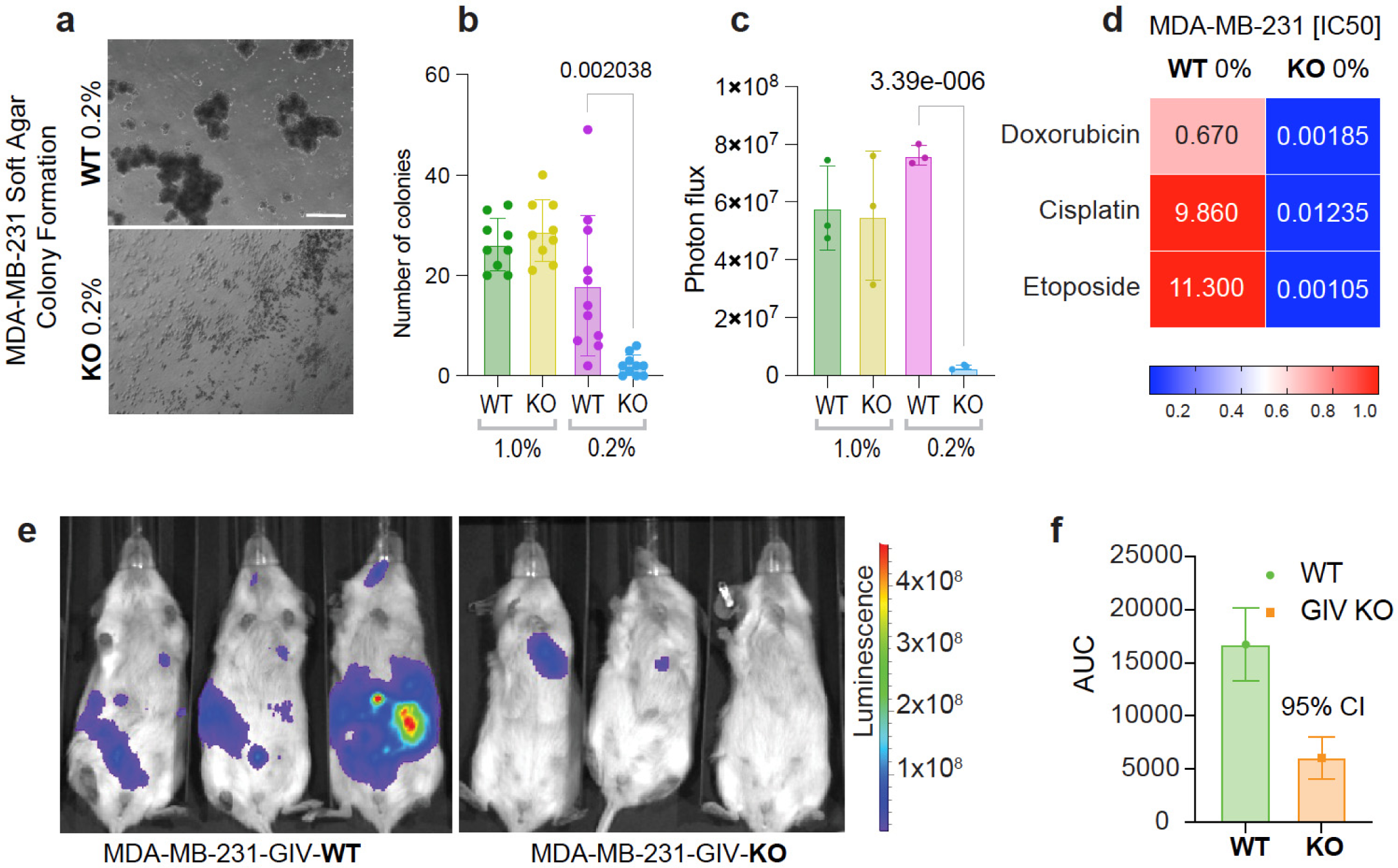
GIV-dependent autonomy is required for growth in growth factor-restricted conditions, esilience to chemotherapeutics and metastatic seeding. **a-c.** Images (a) display representative fields acquired by light microscopy from soft-agar colony growth assays see Methods) on luciferase-expressing parental (WT) and GIV-KO MDA-MB-231 cells, conducted in serum-estricted conditions. Scale bar = 30 µm. Bar plots (b) display the number of colonies/high-power field (HPF) of parental (WT) and GIV-KO MDA-MB-231 cells, quantified using images in 1% and 0.2% FBS (0% FBS for 1 week prevents colony formation altogether). Bar graphs (c) display the average photon flux emitted from colonies in above explained experimental setup. Results in b-c are representative of 3 biological repeats, and 20-40 HPFs were quantified in each repeat. Error bars in c represent S.E.M. p values were calculated by two-ailed t-test. **d**. IC50 values for three commonly used chemotheraputic drugs, displayed as a heatmap. **e-f.** Representative whole mouse bioluminescence images (e) of mice acquired on day #22 after intracardiac njection of serum-deprived parental (WT) and GIV-KO MDA-MB-231 cells in panel a. The logarithmic pseudo-color scale depicts the range of bioluminescence values with red being the highest and blue the lowest. Bar graphs (f) display mean values ± S.E.M. for area-under-the-curve (AUC) for total bioluminescence from days 1-29 after intracardiac injection for each group. N = 5 mice per condition. See also **Figure S3** for similar studies on MCF7 cells (that lack endogenous GIV).

Previous work has documented the absence of full-length GIV in MCF7 cells, the most widely used ER-positive breast cancer cell line (43.6% of total PubMed citations(32)). These MCF7 cells depend on the growth hormone estrogen to proliferate. We found that restoring GIV expression in these cells using a Tol2-based transposon vector was sufficient to enable estrogen-independent growth, as determined using the ER-antagonist Fulvestrant (**Fig S3a-b**) and Tamoxifen **Fig S3b**). GIV was also sufficient for the growth of MCF7 as spheroids in soft agar under estrogen- and serum-restricted conditions (0.2%; **Fig S3c-d**). This impact of GIV on cell growth/survival in serum-restricted conditions was limited to growth factors and hormone, but not for the CDK4/6 inhibitor, Palbociclib (**Fig S3b**), a commonly used therapeutic agent in metastatic ER+ breast cancers, which blocks the cell cycle transition from G1 to S by inhibiting the kinase activity of the CDK/cyclin complex. Findings suggest that GIV-dependent growth signaling autonomy may be sufficient for growth factor and hormone-restricted growth.

### ‘Autonomy’ represents a distinct cell state that is self-sufficient in EGFR/ErbB growth signaling

RNA sequencing studies revealed a set of 30 genes (27 up- and 3 downregulated) and two miRNAs were most differentially expressed (DEGs; Log fold change >5; **Fig 4a**; **Supplemental Information 3**) between the autonomous WT and the GIV-KO cells. These genes were up- and downregulated in WT and KO cells, respectively, in response to serum depravation (**Fig 4b**), a pattern that was strikingly similar to those observed previously for signatures of stemness (**Fig 1g**) and EMP (**Fig 2a-e**). No genes were significantly differentially expressed between the two cell lines when cultured in 10% serum. The list of upregulated DEGs was notable for the presence of EGF (**Fig 4c**); a reactome pathway analysis confirmed that this list was significantly enriched in genes that participated in the EGFR/ErbB-signaling pathway (**Fig 4d**). Pathway analysis of the downregulated DEGs was notable for cellular processes related to the extracellular matrix (ECM), e.g., collagen formation, assembly, and degradation, activation of metalloproteinases and degradation of ECM (**Fig S4**).

**Figure 4.**
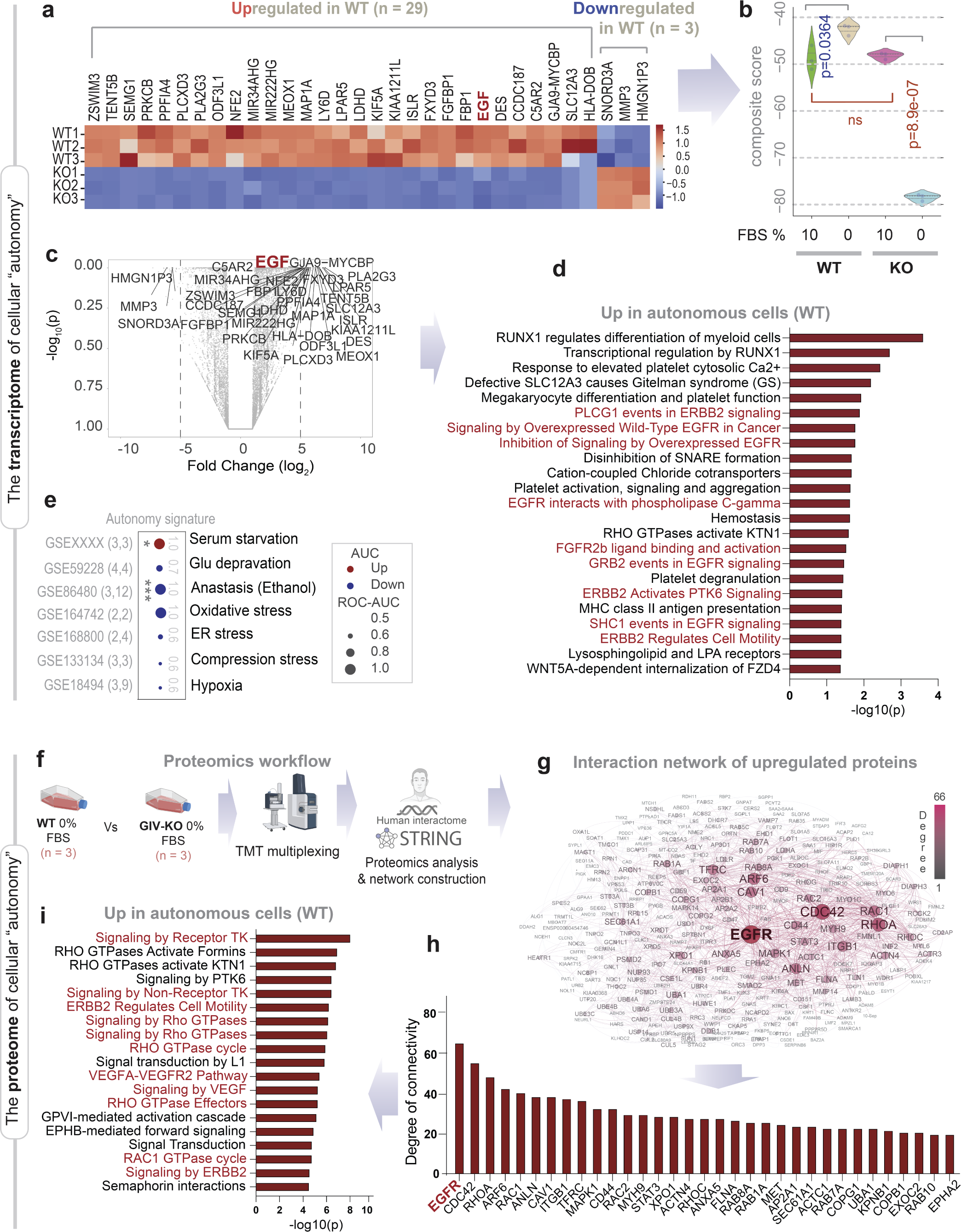
Genes and proteins differentially expressed between autonomy-endowed (WT) and- impaired GIV-KO) MDA-MB-231 cells. **a.** Heatmap displays DEGs (29 up- and 3 down-regulated; LogFC >5, pAdj <0.01) in MDA MB-231 parental (WT) cells compared to its GIV-depleted (KO by CRISPR) counterparts grown in 0% FBS for 16 h. **b.** Violin plots display the composite score of the DEGs in a (used as an ‘autonomy signature’) in parental (WT) and GIV-depleted (GIV-KO) MDA MB-231 cells. **c.** Volcano plot of the differentially expressed genes between WT and GIV-KO MDA MB-321 cells. **d.** Reactome pathway analyses of the pathways enriched in the genes upregulated in WT vs KO MDA MB-231 cells. Red, pathways associated with EGFR/ERBB signaling. See also **Figure S4** for the Reactome pathway for he genes down-regulated in autonomy-endowed (WT) cells. **e.** Expression of autonomy signature in cells subjected to various stressors is visualized as bubble plots of ROC-AUC values (radius of circles are based on the ROC-AUC) demonstrating the direction of gene regulation (Up, ed; Down, blue) for the classification of control vs stress conditions. MDA MB-231 cells were used in all except anastasis, which was done in HeLa cells. *p* values based on Welch’s T-test (of composite score of gene expression values) are provided either as exact values or using standard code (‘’p>0.1, ‘.’p<=0.1, *p<=0.05, *p<=0.01, ***p<=0.001) next to the ROC-AUC. See **Figure S5a-j** for visualization of the data as violin plots. **f-h**. Schematic (f) of the workflow for proteomic analyses. Briefly, differentially expressed proteins (upregulated n WT, compared to KO; n = 3 samples each), as determined by TMT proteomics were used for protein-protein network construction using the human interactome database curated by STRING (Search Tool for the Retrieval of Interacting Genes/Proteins; https://string-db.org/). The resultant network of protein-protein interaction is displayed in g. Node size indicates the degree of connectivity within the network. Degree of connectivity of nodes with a z score of degree (Z_d_) >=1 is represented as a bar plot in h. i. Reactome pathway analyses of the most connected nodes (Z_d_ >= 3) that are upregulated in WT vs KO MDA MB-231 cells. Red, pathways associated with EGFR/ERBB signaling.

Using the composite score of expression of the 30 genes as a signature of growth signaling autonomy (henceforth, “Autonomy signature”) we navigated a wide range of cellular stress response states to assess specificity to growth-factor deprivation (as opposed to non-specific ‘stress response’). The autonomy signature was induced in MDA MB-231 cells challenged with serum deprivation (**Fig S5c**) but not glucose deprivation, oxidative stress, hypoxia, ER-stress, or mechanical compression (**Fig 4e; Fig S5c-i**). The signature was suppressed in HeLa cells undergoing anastasis, a process of resurrection in which cancer cells can revive after ethanol-induced apoptosis (**Fig S5a-b; Fig 4e**). Furthermore, the autonomy signature had no overlaps with other signatures that are either approved for clinical use by the FDA or in clinical trial for their utility in the management of breast cancers (**Fig S5j**). Together, these findings indicate that the gene set of growth signaling autonomy are unique; they represent a distinct ‘cellular state’ that is induced under serum-restricted conditions and requires GIV.

Tandem Mass Tag (TMT)-based quantitative proteomics studies (**Fig 4f**) revealed that that autonomous WT cells differential express a distinct set of proteins, which included EGFR (2.308-fold; **Supplemental Information 4**). When we constructed a protein-protein interaction (PPI) network using this list of upregulated proteins fetched from the human interactome (see network edges in **Supplemental Information 5**), EGFR emerged as the node with highest degree of connectivity (**Fig 4g-h; Supplemental Information 6**). The most connected proteins (i.e., nodes of the PPI network with Z_d_ >= 3) that are upregulated in autonomous WT cells were, once again, found to be significantly enriched in proteins that participate in the EGFR/ErbB2 signaling pathway (**Fig 4i**).

Thus, the transcriptomic and proteomic studies agree; both reveal an upregulated EGFR/ErbB2 signaling pathway in the autonomous WT cells during serum-restricted conditions and confirm the requirement of GIV in such upregulation. Intriguingly, both *EGF* gene and EGFR protein emerged from these ‘omics’ studies, with an enrichment of its immediate downstream signaling (e.g., signaling via Grb2, PLCγ, CDC42, Rho and Rac GTPases), and both secretory and endocytic trafficking proteins (e.g., COP1, RABs, SNARE, EXOC1 ARF6, AP2-subunits, CAV1 proteins; **Fig 4h**). Findings show that the transcriptome and proteome of growth signaling autonomy supports both auto-/paracrine secretion and signaling within the EGFR/ErbB pathway. They also provide a gene signature for growth signaling autonomy, which is exclusively induced on-demand during scarcity of resources.

### Growth signaling autonomy is induced in CTCs

Next, using the newly derived autonomy signature as a computational tool, we sought to navigate the various steps within the cancer initiation and progression cascade. A microarray dataset generated using xenografts of MDA MB-231 cells implanted into inguinal and axillary fat pads of NOD scid female mice, which included samples representing all major steps of the cascade was prioritized (see **Fig 5a***-left*). To our surprise, the autonomy signature was neither induced in primary tumors, nor in metastases; it was induced exclusively in CTC isolated from blood samples (**Fig 5a***-right*). Findings in mice were conserved in humans; the autonomy signature was induced in human CTCs (**Fig 5b**) but not in human primary tumors (when compared to normal, across all molecular subtypes; **Fig S5k**). When primary tumors from patients with detectable CTCs were compared to those without, the autonomy signature was higher in tumors that shed CTCs (**Fig 5c**). Although this degree of specificity (for CTCs) was surprising, that the autonomy signature was induced in CTCs is consistent with the near total lack of biologically active EGF in serum (see *Introduction*).

**Figure 5.**
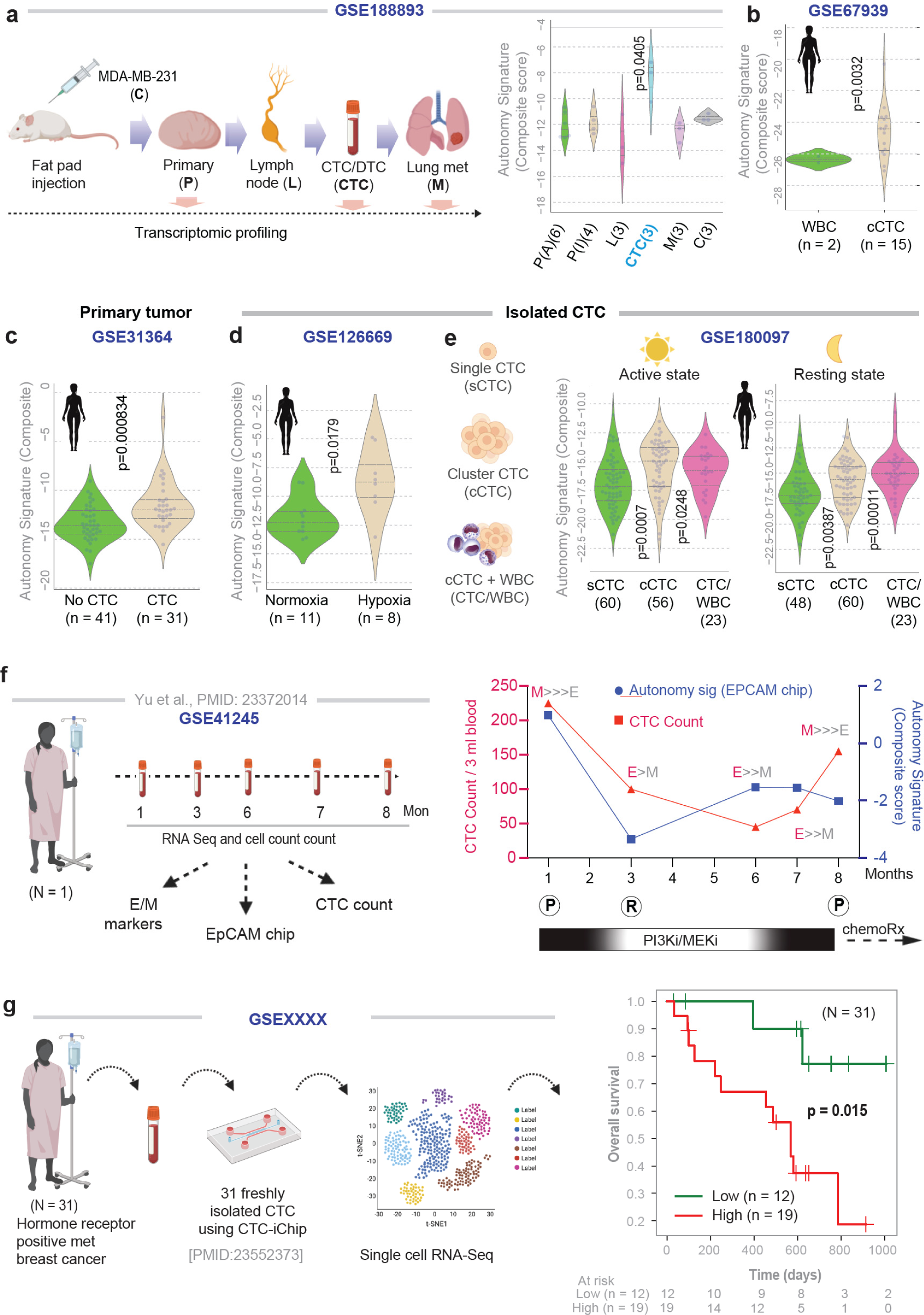
Autonomy signature is induced in CTCs, tracks treatment response and carries poor prognosis. **a.** *<u>Left</u>:* Schematic displays of the study design for a publicly available dataset, <u>GSE103639</u>, in which the MDA MB-231 breast cancer cell line was injected into NOD scid female mice and then RNA seq was carried out for primary tumors from both axillary (P-A) and inguinal (P-I) locations, lymph nodes (L), isolated CTCs, and metastases to the lung (M). *<u>Right</u>:* Violin plots display the composite score for autonomy signature in all samples and assessed for statistically significant differences compared to the primary tumor by Welch’s t-test. **b-e**. Violin plots display the composite score for autonomy signature in various human datasets comparing 15 CTC clusters from a single patient during one blood draw against WBC control (b), primary tumors with/without detectable CTCs (c), CTCs challenged or not with hypoxia (d) and single vs. clustered CTCs collected during different times of the day (e). Statistical significance was assessed Welch’s t-test. **f.** *<u>Left</u>*: Schematic displays study design used in a publicly available dataset, <u>GSE41245</u>. *<u>Right</u>*: Graph tracks the abundance of CTCs (CTC count; blue, left y axis), overlaid on composite score of the levels of expression of genes in the autonomy signature in CTCs (EpCAM chip; red, right y axis). The ratio of epithelial (E) vs mesenchymal (M) markers, as determined by qPCR and reported in the original study by RTqPCR on a panel of markers is indicated. X-axis (time) is annotated with the therapeutic regimen and clinically determined disease status; (P) progression, and (R), response. **g.** *<u>Left:</u>* Schematic showing the study design in which CTCs were isolated from 31 unique subjects with MBC using CTC-iChip microfluidic device and underwent single-cell CTC RNA-seq (<u>GSE215886</u>). *<u>Right:</u>* KM curves of overall survival on the same cohort, stratified based on high (red) vs low (green) autonomy signature. Statistical significance (p value) was determined by the log-rank test.

Next, we interrogated diverse human CTC datasets that were generated by independent groups, in which the metastatic proficiency of the CTCs was experimentally validated using xenograft models. For example, intratumoral hypoxia is a known driver of intravasation of clustered CTCs with high metastatic proficiency (18). We found the autonomy signature significantly higher in hypoxic clusters of live CTCs were compared against their normoxic counterparts (**Fig 5d**), all drawn from a breast cancer patient and labeled with HypoxiaRed, a cell-permeable dye that tags hypoxic cells based on their nitroreductase activity (18). The shedding of CTCs is known to peak at the onset of night (41), when they display more metastatic proficiency (19). It is also known that compared to single CTCs, the metastatic proficiency of CTC clusters (15) and CTC-WBC clusters (8) are higher. The autonomy signature was induced in CTC/CTC-WBC clusters (compared to single CTCs; **Fig 5e**); the significance of such induction was higher in CTC-WBC clusters that were collected at night (**Fig 5e**).

### Autonomy signature in CTCs tracks therapeutic response and prognosticates outcome

CTCs exhibit dynamic changes in abundance and epithelial and mesenchymal composition during treatment (10); we asked if/how treatment might impact the autonomy signature. We analyzed a dataset comprised of CTCs serially collected during a previously published study on an index patient (**Fig 5f***-left*), which displayed reversible shifts between these compositions accompanying each cycle of response to therapy (R) and disease progression (P). The autonomy signature was rapidly downregulated (alongside CTC count) during treatment initiation, which coincided with therapeutic response (1-3 mo.; **Fig 5f***-right*). The signature was subsequently induced from 3^rd^-7^th^ month (despite continuation of treatment and low CTC counts) and preceded clinically confirmed disease progression at the 8^th^ mo. which necessitated salvage chemotherapy (**Fig 5f***-right*). The signature did not show any discernible relationship with the relative amounts of E/M compositions, which is in keeping with our prior observation that autonomy is associated with EM-plasticity (**Fig 2**).

We next re-analyzed a single-cell RNA seq dataset generated using 135 viable CTC samples, from 31 unique patients with hormone receptor positive MBC, for whom follow-up and outcome data were available (overall survival, as updated on 2/10/2020 (7)). CTCs were freshly isolated directly from whole blood using a CTC-iChip microfluidic device (42) (**Fig 5g***-left*). A Kaplan-Meier survival analysis revealed a higher 3-year mortality risk among those with high autonomy signature in CTCs compared to those with low expression of the same (*p*=0.015; **Fig 5g***-right*).

### Autonomy is associated with the potential to re-epithelialize, evade the immune system, and proliferate

CTCs must display plasticity between epithelial and mesenchymal states to complete the metastatic process (10), and the EGF/EGFR pathway has been identified as 1 of the 14 major pathways that support EM-plasticity (39). We asked if the self-sustained EGF/EGFR signaling program in autonomy is specifically associated with ‘reversibility’ of the EMT process. We took advantage of a dataset in which Her2-transformed human mammary epithelial (HMLE) cells were either programmed for reversible EMT (induced by TGFβ) or to a stable mesenchymal phenotype (by chronic exposure to the ErbB inhibitor, lapatinib) (**Fig 6a**). Xenograft studies using these programmed cells had confirmed that reversible, but not stable mesenchymal phenotype produces long-bone metastases (43). We found the autonomy signature to be higher in cells programmed for reversible EMT compared to both stable mesenchymal cells and established bone metastases (**Fig 6b**). This indicates that the autonomous state is present in transformed cells that carry the potential to undergo dynamic transitions (EM-plasticity) but is lost when cells get stuck in either stable mesenchymal (as during the emergence of resistance to Lapatinib) or re-epithelialized states (as in established metastatic colonies).

**Figure 6.**
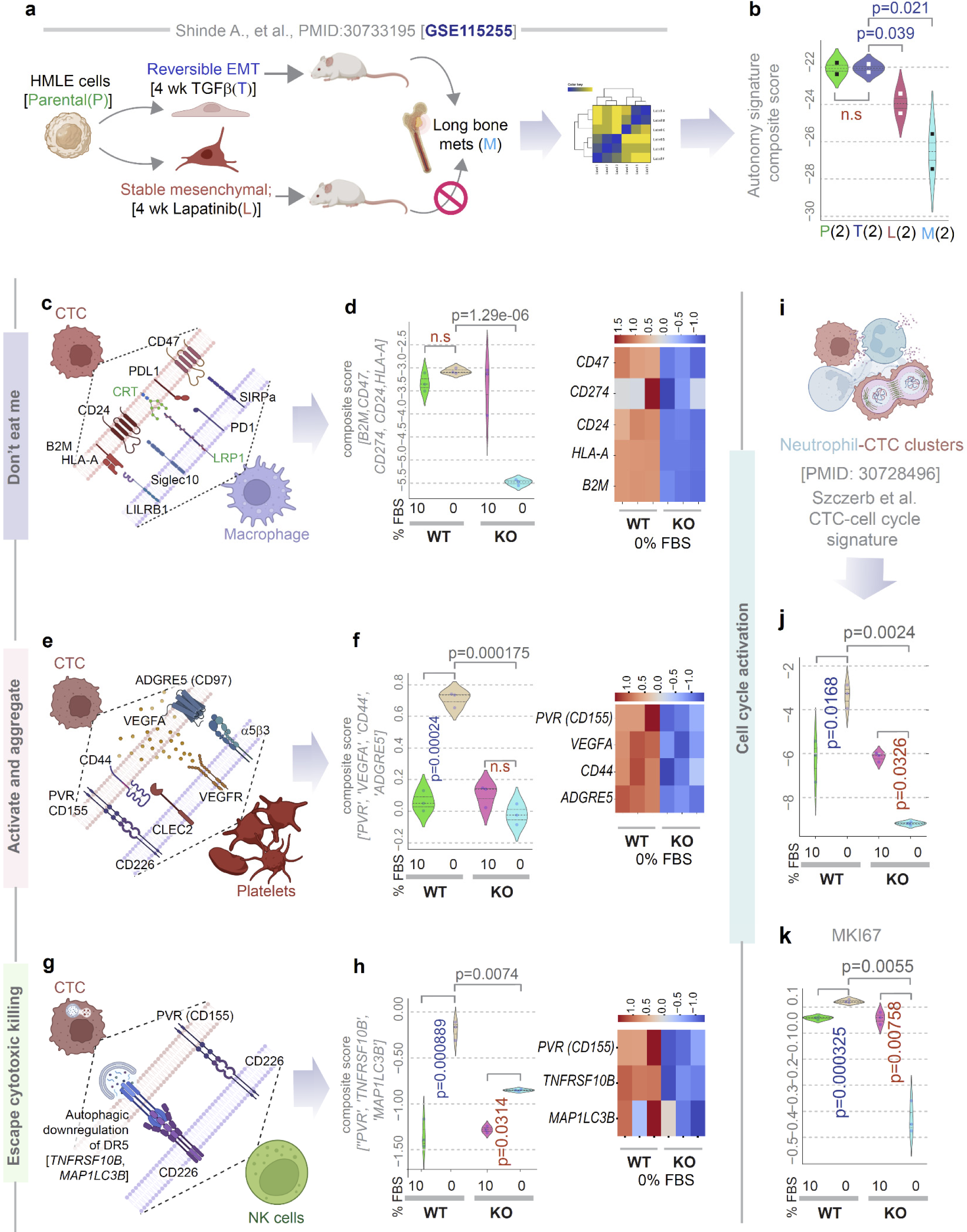
Autonomy is associated with reversible EMT, capacity to evade immune cells and proliferate. **a-b.** Schematic (a) displays the key steps in study design using Her2-transformed mammary epithelial cells HMLE, parental). Induction of reversible EMT (by TGFβ) but not stable EMT (by Lapatinib) is associated with he development of long-distance metastases to bones (BM). Parental, reversible EMT, stable mesenchymal and bone metastases-derived clones of HMLE cells were subjected to RNA seq. Violin plots (b) display the composite score of the genes in the autonomy signature in the HMLE clones. *p* values based on Welch’s T-test, comparing TGFβ-induced reversible EMT and other clones. **c-h.** Schematics summarize the paired CTC-macrophage (c), or CTC-platelets (e) or CTC-NK cell (g) components known to enhance the metastatic potential of heterotypic CTC clusters(45). Violin plots (left panels; d, f, h) show the composite score of various markers of immune evasion in CTCs in parental (WT) and GIV-depleted (GIV-KO) MDA MB-231 cells. Heatmaps (right panels; d, f, h) show the expression of each gene. See **Figure S6** for violin plots for the individual genes. i**-k**. Schematic (i) summarizes the study showing cell cycle activation in CTC-neutrophil clusters. Violin plots show either the composite score of expression of genes in that specific CTC-associated cell cycle signature (j) or levels of expression of *MKI67* (k) in parental (WT) and GIV-depleted (GIV-KO) MDA MB-231 cells. *Statistics*: *p* values based on Welch’s T-test. Blue and red font for p values indicate, significant up- or downregulation, respectively.

EMT and stemness in tumor cells correlate with immune checkpoint expression and complex interactions with platelet and immune cells (35, 44); similarly, the metastatic proficiency of the CTCs is regulated by interactions with the platelets and immune cells (45). We found that all major CTC markers that are known to be critical for the assembly of the CTC-monocyte (**Fig 6c-d**), the CTC-platelet (**Fig 6e-f; Fig S6a-f**) and the CTC-NK cell (**Fig 6g-h**) synapses were expressed at significantly higher levels in the autonomous WT cells compared to their GIV-KO counterparts exclusively in 0% FBS conditions. These findings suggest that autonomous WT, but not the autonomy-impaired GIV-KO cells are likely to be able to mount an immune evasion [“do not eat me”] response by escaping phagocytosis by monocytes, triggering platelet aggregation and activation which shields CTCs from NK cells, and finally, evading cytolytic killing by NK cells.

Besides immune evasion, CTC-neutrophil interactions are known to induce the expression of CTC genes that outline cell cycle progression, leading to more efficient metastasis formation (45). This neutrophil-related pro-proliferative signature was highly expressed in the autonomous WT cells but suppressed in the autonomy-impaired GIV-KO cells upon serum deprivation (**Fig 6i-j**). A similar pattern was seen also for the universal proliferation marker gene, MKI67 (**Fig 6k**), and two other gene sets [Gene Set Enrichment Analysis (GSEA)] for cell cycle progression, KEGG and BIOCARTA (**Fig S6g-h**).

These findings suggest that the autonomous state in serum-restricted condition is associated with three key CTC properties that are essential for metastasis, i.e., plasticity, immune evasion, and proliferative potential.

## Discussion

In this work, we validate a model system for studying one of the hallmarks of cancers, i.e., self-sufficiency in growth signaling or growth signaling autonomy, define the transcriptome, proteome, and phenome of such autonomous state, and unravel its role during cancer progression. Findings show that the autonomous state is prominently induced in a subset of CTCs and empower them with key properties which may make them better ‘seeds’ for metastasis (see **Fig 7**). Elaborated below are the three major implications of these findings.

**Figure 7.**
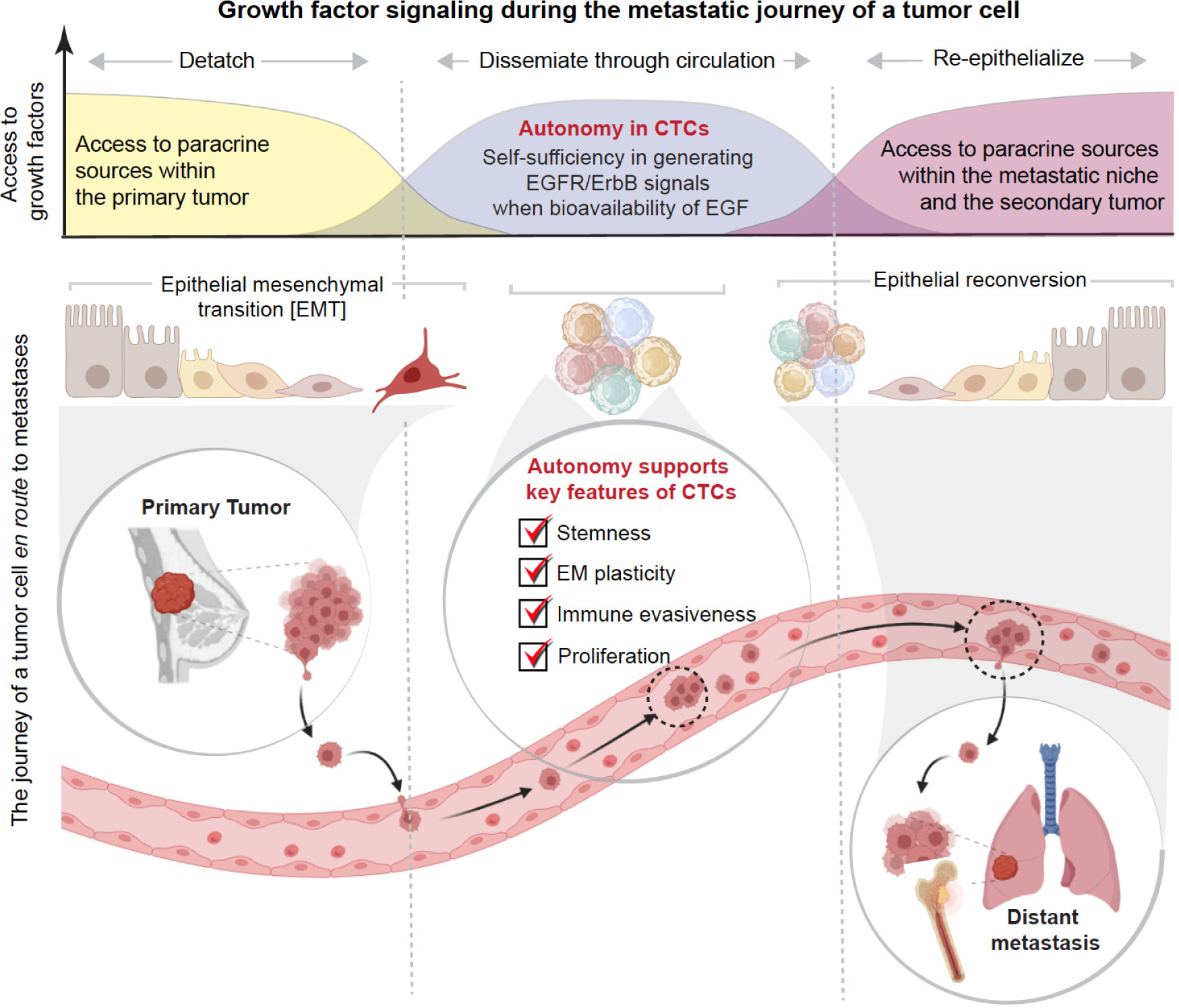
Summary of findings and working model. Schematic showing the various phases of the journey of umor cells *en route* to seeding metastases. *<u>Top</u>*: Growth factor signaling and source during that journey. *<u>Bottom</u>*: Epithelial to mesenchymal (EMT) and mesenchymal to epithelial (MET) transition states in primary (left) or metastatic (right) tumors and the transitioning state in CTC clusters. Growth signaling autonomy, which is seen n CTCs, appears to support a self-sufficient EGFR/ERBB signaling program that is required for re-epithelialization during metastasis. Other properties of autonomous CTCs include, stemness, EM-plasticity, mmune evasiveness, and cell proliferation.

### The autonomous state embodies key features of CTCs that confer metastatic potential

We found that the autonomous state that is unique to a subset of CTCs supports a gene expression program for stemness (i.e., oncogenic de-differentiation), proliferation, immune evasiveness, and EM-plasticity. Of these features, the most distinct one is EMP, because much like what we see in the case of growth signaling autonomy, EMP is unique to CTCs and very rare in primary tumor cells or in metastases (46). Autonomy is reduced/lost in cells that are unable to transition between E↔M states. These findings are in keeping with the gathering notion that CTCs are the metastatic precursors that simultaneously express epithelial and mesenchymal markers and best display dynamic E→M and M→E transitions (10). Such a hybrid state in which CTCs have acquired only a partial mesenchymal state allows rapid E→M transitions needed to migrate and intravasate, and quick M→E reversals to reinitiate a tumor at a distant site (47). Consequently, epithelial-type CTCs with a restricted mesenchymal transition initiate metastases efficiently, whereas mesenchymal-type CTCs don’t (12). Consistent with prior findings that CTC clusters express cell-cell adhesion proteins that are components of tight junctions and desmosomes (15, 48), we show that the autonomous state is endowed with gene expression program to support tight junctions (which are essential for the formation of CTC clusters). Divergent from classical EMT, EM-plasticity is known to also induce unique immunomodulatory effects (49, 50); consistent with this notion, our model was also accompanied by a diverse array of immune-evasion machinery. Because EM-plasticity has been broadly implicated in metastasis, chemoresistance and immunosuppression (51), and is present in autonomy-endowed cells that outperformed autonomy-impaired cells in their ability to initiate metastases in mice, we conclude that autonomy-endowed CTCs are likely to be more efficient in seeding metastases than those CTCs that are autonomy-impaired.

Mechanistically, growth signaling autonomy is supported by a secretion-coupled sensing circuit at the Golgi apparatus (29, 30) which controls the secretory flux and organelle shape (compact vs fragmented stacks when the circuit is enabled or disabled, respectively). These findings are in keeping with prior work showing that CTC population level behavior is dominated by a few high-secreting cells (52), and that Golgi shape (fragmentation vs compact) is a strong correlate of CTC-EMT and invasiveness (53). Although it remains unknown if/how the Golgi-resident circuit begets EM-plasticity, we conclude that growth signaling autonomy and phenotypic plasticity, two hallmarks of cancer, co-exist in CTCs.

### Autonomous CTCs support a self-sufficient EGFR/ErbB signaling program

Our transcriptomic and proteomic analyses pinpointed with surprising convergence that the autonomous state supports a self-sufficient EGFR/ErbB-centric signaling program in the absence of external growth factors. While most cells can be stimulated by growth factors made by neighboring cells—via the process of paracrine signaling—many cancer cells acquire self-sufficiency such that they sense and respond to what they synthesize and secrete, creating a positive feedback sense-and-secrete loop known as autocrine stimulation (54). Such a type of self-sufficiency or autonomy in a cancer cell obviates its need to depend exclusively on the surroundings, especially during the intravascular journey (when CTCs cannot access biologically active EGF) and during the initial phase of avascular growth of a CTC that has just extravasated to a new site. Although examples of such autonomy exist in the case of PDGF and TGFα by glioblastomas and sarcomas, respectively, and MDA MB-231 cells are known to secrete EGF(55), autocrine autonomy in the EGF/EGFR pathway has not been described previously and mechanisms that support such sense-and-secrete loops in eukaryotes had remained elusive. By demonstrating that a secretion-coupled-sensing machinery at the Golgi (29, 30) that requires scaffolding by GIV is essential for tumor cells to achieve a state of self-sufficiency in EGFR/ErbB signaling, it is not unusual for us to find that such a state is endowed with a multitude of pro-oncogenic biological processes that are known to be supported by EGFR/ErbB signals, including cell anchorage-independent cell growth, stemness, EM-plasticity and metastasis (56). Findings are also in keeping with prior observation of *EGFR* gene induction in EMT that is accompanied by plasticity and tumorigenicity (but not in EMT alone (57)). Although it remains unclear if our findings are related to the previously reported prognostic roles of high EGFR (58, 59) and Her2 (60) in the serum of patients with breast cancers, EGFR/ErbB2 has been detected in CTCs consistently during serial blood draws (61) and was found to be activated (as determined by the presence of its phosphorylated state (62) in CTCs) with increasing frequency often during MBC progression (62). Because inhibition of EGFR prevents CTC clustering and diminishes metastatic potential (63), it is possible that higher autonomy observed in the CTC-clusters (compared to single-CTCs) requires the autonomous signaling machinery (the sense-and-secrete loop) for CTCs to remain clustered and maintain metastatic potential. It is possible that the EGF-predominant autocrine loop maintains CTC-junctions within clusters by triggering both the secretion of junctional proteins/complexes (e.g., E-cadherin) from the Golgi to the plasma membrane (PM) (64) and their subsequent activation at the junctions (65).

### Autonomy signature could identify CTCs that are the ‘fittest’ precursors to metastases

In the absence of available tools to objectively assess, report and track the metastatic potential of CTCs, their use in prognosticating risk of relapse or to track the emergence of therapeutic resistance in real-time and adapt the clinical response has yet to be realized. We show that a gene signature for growth signaling autonomy can track treatment response in a single index patient, and more importantly, the signature prognosticates outcome in a dataset of 31 unique patients. Molecular markers of the metastatic potential of CTCs have been described before, e.g., RPL15 (7) and a 17-gene signature (66). While RPL15 was identified during an *in vivo* genome-wide CRISPR activation screen to identify genes in breast cancer patient-derived CTCs that promote their distant metastasis in mice, the latter was trained on a cohort of normal vs primary tumor and whole blood from patients. Unlike both these instances, in which investigative approaches were geared to identify CTC-specific markers, we stumbled upon the autonomy signature as a portal into the metastatic proclivity of CTCs by serendipity, using the gene expression signature for autonomy. Regardless, of how it was identified, our findings not only add growth signaling autonomy to the growing list of the parameters that help define the ‘fitness’ of CTCs, but also provide a methodology to objectively measure (using a gene signature) the degree of autonomy (and hence, the fitness) of CTCs to serve as ‘seeds’ for metastases. Prospective studies are required to further investigate the clinical utility of this signature.

In conclusion, our work provides insights into what determines the success of a CTC to serve as metastatic “seed”. By demonstrating that the detached tumor cells (*sans* ECM contact) gain autocrine self-sufficiency in growth factor signaling and phenotypic plasticity in circulation, while maintaining the properties of stemness, proliferation and immune evasiveness (which are seen also in primary tumors), we show the coexistence of two hallmarks of cancer that are relatively unique to CTCs and are intricately intertwined.

## Materials and Methods

### EXPERIMENTAL METHODS

#### Cell lines

MDA-MB-231 cells were grown at 37°C in their suitable media, according to their supplier instructions, supplemented with 10% FBS, 100 U/ml penicillin, 100 μg/ml streptomycin, 1% L-glutamine, and 5% CO2.GIV knock-out (KO) cell lines were generated using Pooled guide RNA plasmids (commercially obtained from Santa Cruz Biotechnology; Cat# sc-402236-KO-2), as described earlier(67). Briefly, these CRISPR/Cas9 KO plasmids consist of GFP and Girdin-specific 20 nt guide RNA sequences derived from the GeCKO (v2) library and target human Girdin exons 6 and 7. Plasmids were transfected into Hela and MDA-MB-231 cells using PEI. Cells were sorted into individual wells using a cell sorter based on GFP expression. To identify cell clones harboring mutations in gene coding sequence, genomic DNA was extracted using 50 mM NaOH and boiling at 95°C for 60mins. After extraction, pH was neutralized by the addition of 10% volume 1.0 M Tris-pH 8.0. The crude genomic extract was then used in PCR reactions with primers flanking the targeted site. Amplicons were analyzed for insertions/deletions (indels) using a TBE-PAGE gel. Indel sequence was determined by cloning amplicons into a TOPO-TA cloning vector (Invitrogen) following the manufacturer’s protocol. These cell lines were characterized in earlier work (29, 68), and modified here for luciferase expression by transducing cells with a lentiviral vector expressing click beetle green luciferase (CBG)(69). We selected a population of cells stably expressing CBG based on resistance to blasticidin expressed in the lentiviral vector as described previously.

MCF7 cells were purchased from the ATCC and verified cells by STR profiling through the University of Michigan Advanced Genomics Core. We cultured MCF7 cells in DMEM medium with 10% serum, 1% penicillin/streptomycin, and 1% glutamax (ThermoFisher Scientific) in an incubator set at 37°C and 5% CO_2_. We stably expressed click beetle green luciferase (CBG) in these cells by lentiviral transduction as described previously(69). To stably express full-length, wild-type GIV in MCF7 and MDA-MB-231-GIV KO cells, we used a Tol2 transposon system. The Tol2 transposon vector uses a CAG promoter to drive constitutive expression of the cDNA for *CCDC88A* (the gene name for GIV) and a hygromycin resistance gene linked by a P2A sequence (Vector Builder). We co-transfected cells with the Tol2 GIV transposon and a Tol2 transposase (Vector Builder) at a 3:1 ratio of micrograms of plasmid DNA using Fugene 6 (Promega) according to the manufacturer’s directions. Control cells underwent transfection with an empty Tol2 transposon and Tol2 transposase at the same ratio of plasmid DNA. Two days after transfection, we added hygromycin (Sigma-Aldrich) to select batch populations of cells stably expressing GIV. After obtaining stable cell lines, we did not culture resultant MCF7-GIV cells in hygromycin. We stably expressed CBG in MCF7 cells as detailed for MDA-MB-231 cells.

#### Immunoblotting

To verify the expression of GIV in MCF7-GIV and MDA-MB-231-GIV KO/WT cells, equal aliquots of whole-cell lysates (prepared using RIPA buffer) were loaded on a 8% SDS PAGE gel, and immunoblotting was carried out for GIV with the mouse monoclonal antibody H-6 (Santa Cruz Biotechnology) as described previously(70). Immunoblots were analyzed using a Gel Doc system (BioRad, Hercules, CA).

#### RNA sequencing and identification of DEGs

WT and GIV-KO MDA-MB231 cells grown in 0% and 10% serum concentration in p10 dishes (Corning) for 16 h prior to harvest, and cell pellets were subsequently processed for RNA extraction using a kit (R2052, Zymo Research) as per manufacturer’s protocol. Isolated RNA has been processed for RNA sequencing in the Illumina NovaSeq 6000 platform. Fastq sequence files have been mapped using the human GRCh38 genome. Log normalized CPM expression files are submitted to <u>GSE215822</u>. A list of DEGs is provided in **Supplemental Information 3**.

#### Tandem Mass Tag™ (TMT) proteomics

WT and GIV-KO MDA-MB231 cells were maintained in 0% and 10% serum concentration in p10 dishes (Corning) for 16 h prior to harvest, and cell pellets were subsequently processed for TMT proteomics using LUMOS Orbitrap-Fusion analyzer. Peptides are identified and mapped using Peaks X Pro pipeline. Intensity ratio of each identified proteins in WT MDA-MB231 Vs GIV-KO MDA-MB231 cells has been identified and selected if the significance score >20. The raw proteomics data has been submitted to ProteomeXchange (<u>PXD037253</u>). A list of differentially expressed proteins is provided in **Supplemental Information 4**.

#### Soft agar growth assays

We performed assays with minor modifications from a published protocol(71). Briefly, we made a base layer of 0.7% agar (Sigma-Aldrich) dissolved in DMEM medium, adding 0.2% serum after the agar solution cooled to ∼37°C before transferring 1 ml per well to 6 well plates. After the base layer solidified at room temperature, we prepared 0.35% low melting agar (Sigma) in DMEM medium, adding the same concentration of serum as in the base layer and 5 × 10^4^ cells per ml when the solution cooled to ∼37°C (n = 3 wells per cell type and serum condition). We immediately transferred 1 ml per well of the low melting point agar/cell solution to each well and cooled the plate briefly at 4°C before placing in a cell culture incubator. We added 0.5 ml fresh DMEM medium with 0.2% serum every 2 days. After one week, we obtained 9 bright-field images per well on an inverted microscope (Olympus IX73 with 20X objective). Immediately after microscopy, we added 150 µg/ml luciferin (Promega) to each well; incubated in a cell culture incubator for 10 minutes; and then acquired a bioluminescence image of total viable cells on an IVIS Lumina (30 second image, large field of view) (Perkin Elmer). A person blinded to experimental conditions enumerated colonies and quantified imaging data (Living Image, Perkin Elmer).

#### Cytotoxicity assays

We performed cytotoxicity assays on MCF7 cells as described previously(72). Briefly, we seeded 7.5 × 10^3^ MCF7 WT or MCF7-GIV cells per well in black wall 96 well plates (ThermoFisher Scientific, catalog number 165305). One day after seeding in normal growth medium, we washed cells once with PBS and then added various concentrations of drugs (tamoxifen, fulvenstrant, or 18albociclib; all purchased from Tocris) in phenol red free DMEM (ThermoFisher Scientific) with 4 mM glucose, 1% serum, 1% penicillin/streptomycin, 1% Glutamax, and 10 nM estrogen (n = 4 wells per cell type and concentration). Three days later, we added 150 µg/ml luciferin per well; incubated in a cell culture incubator for 10 minutes; and then acquired a bioluminescence image of total viable cells on an IVIS Lumina (1 minute image, large field of view). We quantified bioluminescence as radiance per well (LivingImage software) and normalized data for each cell type and drug concentration to vehicle only.

#### Animal studies

The University of Michigan Institutional Care and Use of Animals Committee approved all animal procedures. We used 8-10-week-old female NSG mice originally purchased from The Jackson Laboratory and bred in the colony maintained by the University of Michigan Lab Animal Medicine Program. Prior to mouse experiments, we cultured MDA-MB-231 WT and MDA-MB-231-GIV KO/WT cells in serum-free DMEM with 25 mM glucose overnight. We verified that these cells did not lose viability after overnight culture in serum-free medium relative to medium with 10% serum as determined by cell-based measurements of CBG bioluminescence in an IVIS Lumina (Perkin Elmer). We injected 1 × 10^5^ breast cancer cells per mouse (n = 5 per each cell type), verifying positioning of the 30g needle in the left ventricle by return of pulsatile bright red blood as described(73). We imaged bioluminescence with an IVIS Spectrum (Perkin Elmer) in mice at time points shown in the figure legend and quantified data with Living Image software. For each mouse, we calculated the fold-change in bioluminescence relative to the value obtained one day after injection to normalize for variations in injected amounts of cells. We calculated area-under-the-curve ± SEM for total bioluminescence in each group.

### COMPUTATIONAL METHODS

#### Transcriptomic datasets

All publicly available transcriptomic datasets were downloaded from National Center for Biotechnology Information (NCBI) Gene Expression Omnibus website (GEO)(74–76) or European Molecular Biology Laboratory (EMBL) European Bioinformatics Institute (EMBL-EBI) ArrayExpress website(77). All gene expression datasets (**Supplementary Information 1**) were processed separately using the Hegemon (hierarchical exploration of **gene** expression microarrays on-line) data analysis framework (78–80). We did not combine datasets that belong to two different platforms. See **Supplemental Information 1** for the degree of heterogeneity among samples in the datasets used in this work.

#### StepMiner analysis

*StepMiner* is an algorithm that identifies step-wise transitions using step function in a time-series data(81). *StepMiner* undergoes an adaptive regression scheme to verify the best possible up and down steps based on sum-of-square errors. The steps are placed between time points at the sharpest change between expression levels, which gives us the information about timing of the gene expression-switching event. To fit a step function, the algorithm evaluates all possible steps for each position and computes the average of the values on both sides of a step for the constant segments. An adaptive regression scheme is used that chooses the step positions that minimize the square error with the fitted data. Finally, a regression test statistic is computed as follows:

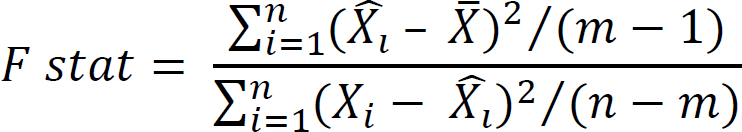

Where *X_i_* for *i* = 1 to *n* are the values, 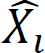 for *i* = 1 to *n* are fitted values. M is the degrees of freedom used for the adaptive regression analysis. 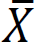 is the average of all the values: 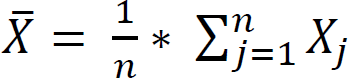. For a step position at k, the fitted values 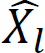 are computed by using 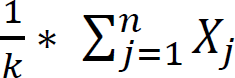 for *i* = 1 to *k* and 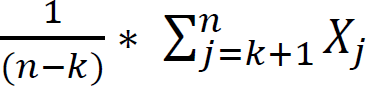 for *i* = *k* + 1 to *n*.

#### Composite gene signature analysis using Boolean Network Explorer (BoNE)

Boolean network explorer (BoNE) provides an integrated platform for the construction, visualization and querying of a gene expression signature underlying a disease or a biological process in three steps: First, the expression levels of all genes in these datasets were converted to binary values (high or low) using the StepMiner algorithm. Second, Gene expression values were normalized according to a modified Z-score approach centered around *StepMiner* threshold (formula = (expr – SThr)/3*stddev). Third, the normalized expression values for every genes were added together to create the final score for the gene signature. The samples were ordered based on the final signature score. Classification of sample categories using this ordering is measured by ROC-AUC (Receiver Operating Characteristics Area Under The Curve) values. Welch’s Two Sample t-test (unpaired, unequal variance (equal_var=False), and unequal sample size) parameters were used to compare the differential signature score in different sample categories. Violin, Swarm and Bubble plots are created using python seaborn package version 0.10.1. Pathway enrichment analyses for genes were carried out via the reactome database and algorithm(82). Violin, Swarm and Bubble plots are created using python seaborn package version 0.10.1. A list of all gene signatures used in this work is provided in **Supplemental Information 2**.

#### Survival outcome analyses

Kaplan-Meier (KM) analyses were done for different gene signatures. The high and low groups were separated based on *StepMiner* threshold on the composite score of the gene expression values. The statistical significance of KM plots was assessed by log rank test. Kaplan-Meier analyses were performed using lifelines python package version 0.14.6.

#### Protein-protein interaction network construction and analysis

Upregulated proteins in WT MDA-MB-231 cells in comparison with the GIV-KO MDA-MB-231 cells are identified with an intensity ratio cutoff of >=2 and with a significance value >= 20. Interaction edges between the identified proteins are fetched from the STRING human interactome database and represented in Figure 4g as a protein-protein interaction network (PPIN) using Gephi 9.02. Degree distribution of the PPIN computed using python network package.

### STATISTICAL ANALYSIS

Gene signature is used to classify sample categories and the performance of the multi-class classification is measured by ROC-AUC (Receiver Operating Characteristics Area Under the Curve) values. A color-coded bar plot is combined with a density plot to visualize the gene signature-based classification. All statistical tests were performed using R version 3.2.3 (2015-12-10). Standard t-tests were performed using python scipy.stats.ttest_ind package (version 0.19.0) with Welch’s Two Sample t-test (unpaired, unequal variance (equal_var=False), and unequal sample size) parameters. Multiple hypothesis correction was performed by adjusting *p* values with statsmodels.stats.multitest.multipletests (fdr_bh: Benjamini/Hochberg principles). Sample number of each analysis is provided with associated plots beside each GSE ID no. or sample name. The statistical significance of KM plots was assessed by log rank test. Pathway enrichment analyses of gene lists were carried out using the Reactome database(83) (http://reactome.org) and the Cytoscape plug-in, CluGo (http://www.ici.upmc.fr/cluego/cluegoDownload.shtml).

## Acknowledgments

We thank Debashis Sahoo (UC San Diego, Boolean Lab) for access to computational resources. This work was supported by the United States National Institutes of Health grants, CA100768, AI141630, CA160911 and CA238042 (to PG), CA238042, CA210152, CA238023, CA225549, CA221807, and CA222563 (to GDL); and CA221807 (KEL). GDL and KEL acknowledge funding from the W.M. Keck Foundation.

## Author Contributions

S.S carried out all the computational analyses in this work. A.F., P.G and G.D.L designed, carried out and analyzed all experiments. S.S and P.G created all the figures. K.E.L. provided new reagents. B.P and K.Y guided S.S and P.G in data analysis and interpretation. S.S and G.D.L wrote methods, and all authors edited the manuscript. P.G and G.D.L conceptualized, designed, and supervised the project. P.G, S.S, G.D.L wrote the manuscript, and all authors edited and approved the same.

