## Supplementary Online Materials for "Growth Signaling Autonomy in Circulating Tumor Cells Aids Metastatic Seeding"

**Affiliations:**

### INVENTORY OF SUPPLEMENTARY MATERIALS

Supplementary Figures and Legends (6)

### SUPPLEMTARY FIGURES AND LEGENDS

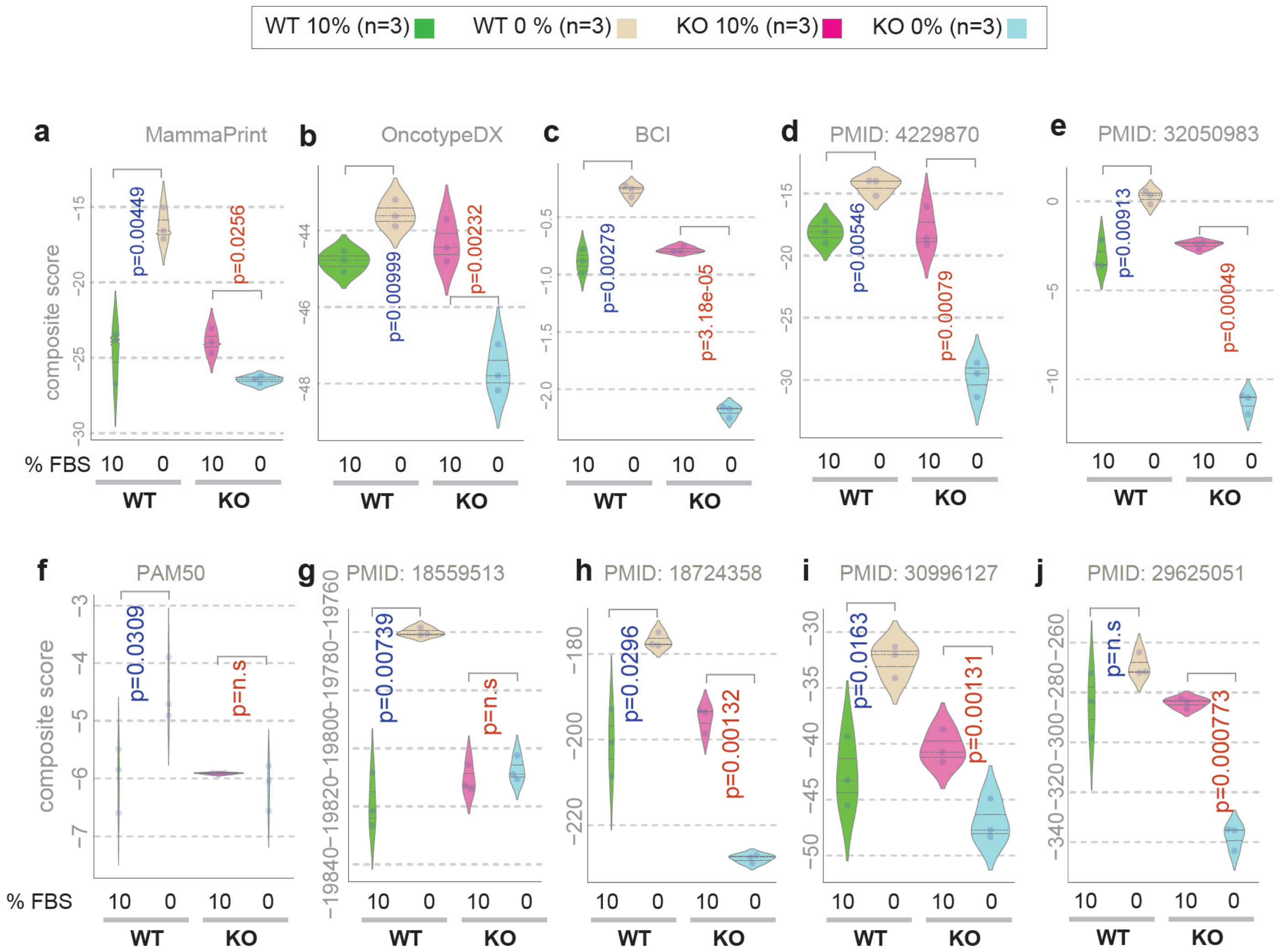


**Figure S1**. [Related to Figure 1]

**GIV-dependent growth signaling autonomy is required for the induction of cancer cell stemness.**

**a-j**. Violin plots display the composite score of selected genes and/or signatures showcased as bubble plots in **Figure 1g**. P values based on Welch's T-test, comparing 10% vs 0% growth conditions in WT (blue) and KO (red) cells. Blue and red font for p values indicate, significant up- or downregulation, respectively.

*Statistics*: *p* values based on Welch's T-test (of composite score of gene expression values) are provided as exact values.

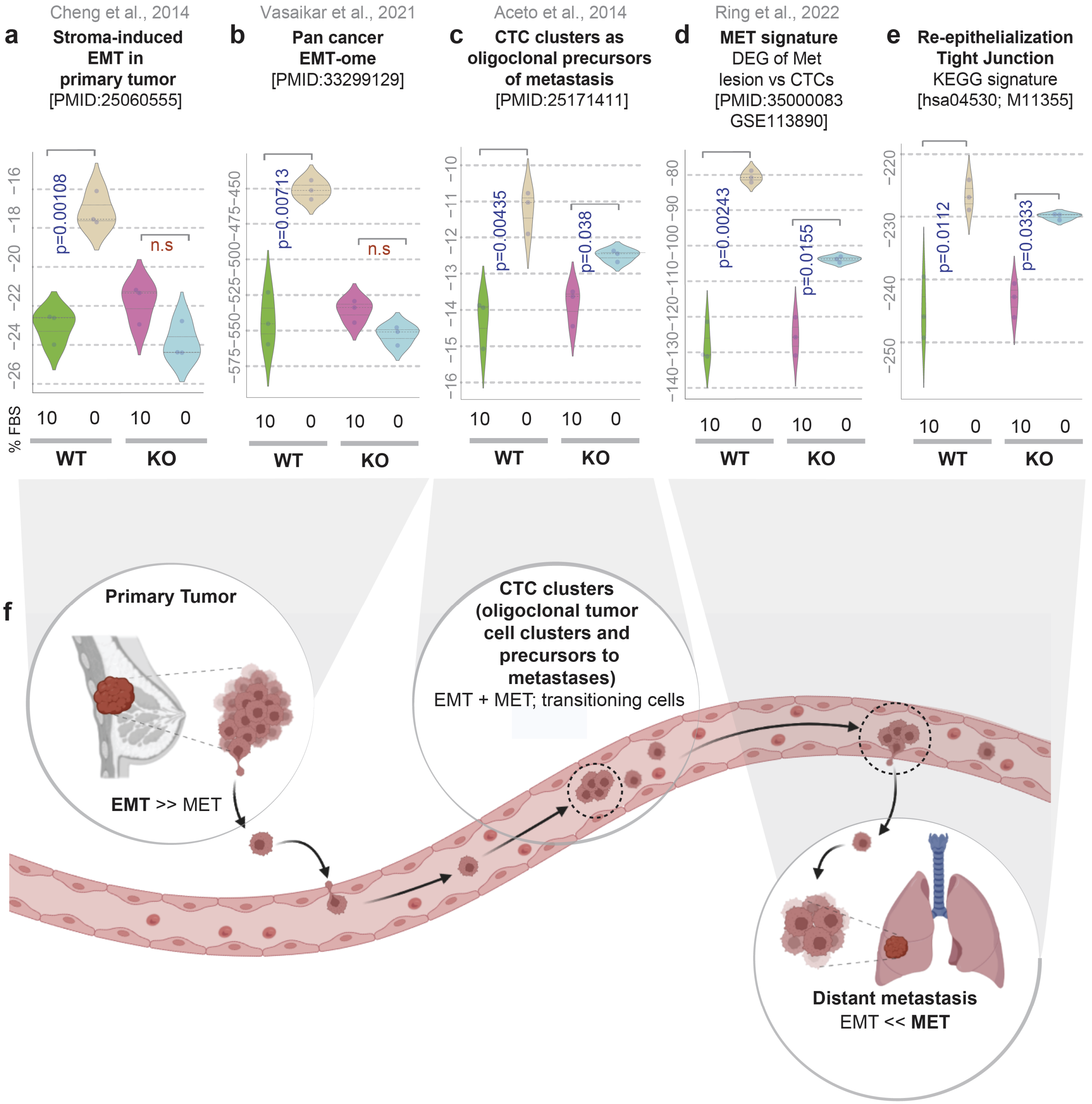


**Figure S2.** [Related to Figure 2]

**GIV-dependent growth signaling autonomy is required for both EMT and MET programs that are essential during distinct steps of tumor metastasis.**

**a-e.** Violin plots display the composite score of various gene signatures in parental (WT) and GIV-depleted (GIV-KO) MDA MB-231 cells, which capture the continuum of epithelial and mesenchymal states that aid in the metastatic spread of breast cancers. <u>*From left to right*</u>: EMT in primary tumors (a-b), to EMT and MET transition capability endowed oligoclonal tumor cell clusters in circulation (CTCs) that have ~23-50-fold increased metastatic potential(*1*) (c), to the MET-predominant process (d) of CTC re-epithelialization associated junctional signatures (e). *p* values based on Welch's T-test, comparing 10% vs 0% growth conditions in WT (blue) and KO (red) cells. Blue and red font for p values indicate, significant up- or downregulation, respectively.

**f.** Summary and working model: Schematic displays the epithelial to mesenchymal (EMT) and mesenchymal to epithelial (MET) transition states in primary (left) or metastatic (right) tumors, and the transitioning state in CTC clusters.

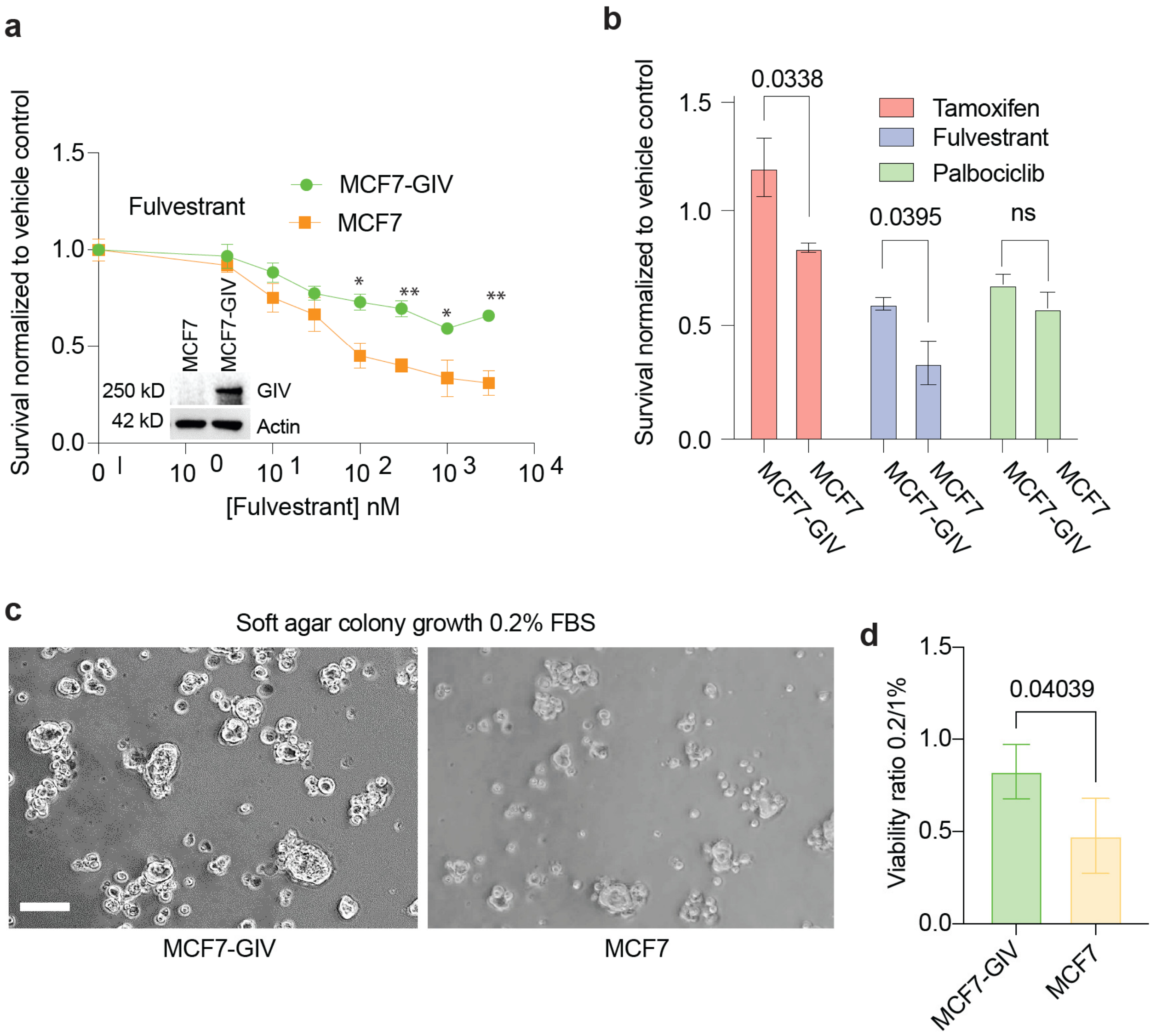


**Figure S3.** [Related to Figure 3]

**GIV is sufficient for therapeutic resistance (to anti-estrogen therapy) and growth factor-restricted proliferation**

**a.** Line graph displays % survival of parental MCF7 cells and clones of the same stably expressing GIV (introduced by Tol2 transposon) after challenge with the indicated concentrations of the selective estrogen receptor blocker (Fulvestrant), as determined by MTT assays. Error bars represent S.E.M (n = 4 biological repeats; each with 3 technical repeats). p values were calculated by one way ANOVA at a given concentration of drug. Only significant values are displayed using standard code (*p<=0.05, **p<=0.01, ***p<=0.001).

**b**. Bar graphs display the proportion of surviving cells (assessed exactly as in g), when challenged with 4 commonly used drugs at $10^2$ nM dose. Error bars represent S.E.M. P values were calculated by one-way ANOVA at a given concentration of drug. Only significant values are displayed.

**c-d**. Images (c) display representative fields acquired by light microscopy from soft-agar colony growth assays (see Methods) on luciferase expressing parental (MCF7) and GIV-expressing (MCF7-GIV) MCF7 cells, conducted in serum-restricted conditions. Scale bar = 30 µm. Bar graph compares the viability ratio of the cells in 0.2% vs 1% serum quantified using the average photon flux emitted from the colonies in c. Error bars represent S.E.M (n = 3 biological repeats; each with 3 technical repeats). p values were calculated by one-tailed t-test.

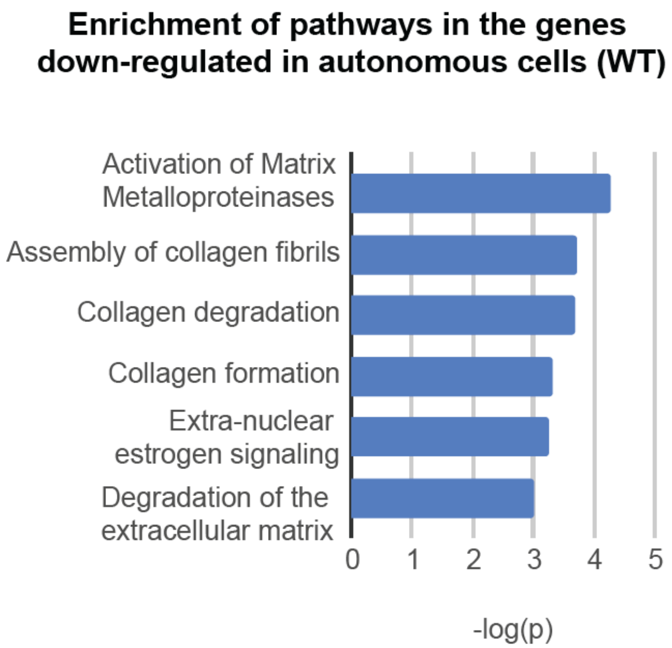


**Figure S4.** [Related to Figure 4]

**Pathway enrichment analysis for differentially upregulated genes in autonomous cells.**

Differentially expressed proteins (downregulated in WT, compared to KO; n = 3 samples each), as determined by RNA Seq were used for pathway enrichment analysis using reactome.

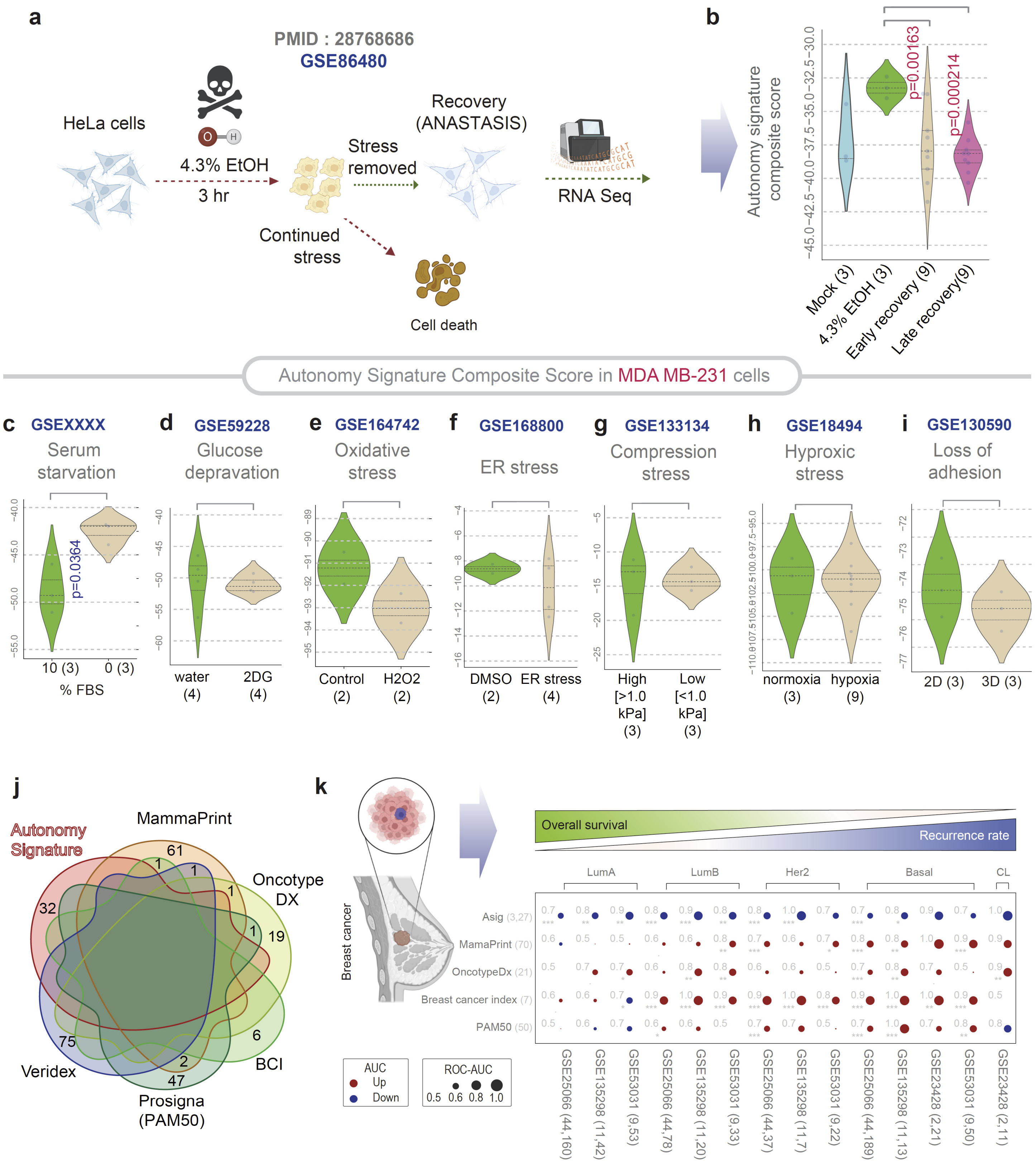


**Figure S5.** [Related to Figure 4]

**Autonomy signature is not a mere stress response; it is relatively specific to growth factor restricted conditions.**

**a**. Schematic (left) summarizes the key steps of experimental design to study gene expression during anastasis of HeLa cells that were challenged with 4.3 % of EtOH followed by recovery (GSE86480). Violin plot (right) shows Violin plots display the composite score of the genes in the autonomy signature in the HeLa cells, collected at various time points during anastasis. Autonomy signature is significantly suppressed early (within 4 h) during recovery and stays suppressed during late (12 h) recovery. *p* values based on Welch's T-test (of composite score of gene expression values) are provided besides each time point compared to the initial condition (prior to EtOH application).

**c-i**. Violin plots show the composite score of the genes in the autonomy signature in MDA MB-231 cells exposed to different forms of stressful growth conditions. *p* values based on Welch's T-test (of composite score of gene expression values) are provided besides each time point compared to the stress-free (initial/control) condition in each case.

**j**. Venn diagram displays the unique and overlapping genes between the newly identified 'Autonomy signature' and five other breast cancer gene signatures with established clinical utility.

**k**. *Top*: Schematic shows the clinicopathological features of various subtypes of breast cancers: (from left to right, Lum A, Lum B, HER2+, Basal and Claudin-low). *Bottom*: The composite score of levels of expression of ASig genes (top row) and genes in other clinically useful signatures (rows 2-5) in the various subtypes of breast cancers compared to normal-like breast tissue is visualized as bubble plots of ROC-AUC values (radii of circles are based on the ROC-AUC) in four independent datasets, demonstrating the direction of gene regulation (Up, red; Down, blue) for the classification of samples (gene signatures in columns; dataset and sample comparison in rows). The direction of gene expression also indicated the abundance of stemlike/dormant cells in primary tumor. p values based on Welch's T-test (of composite score of gene expression values) are provided using standard code (*p<=0.05, **p<=0.01, ***p<=0.001) next to the ROC-AUC.

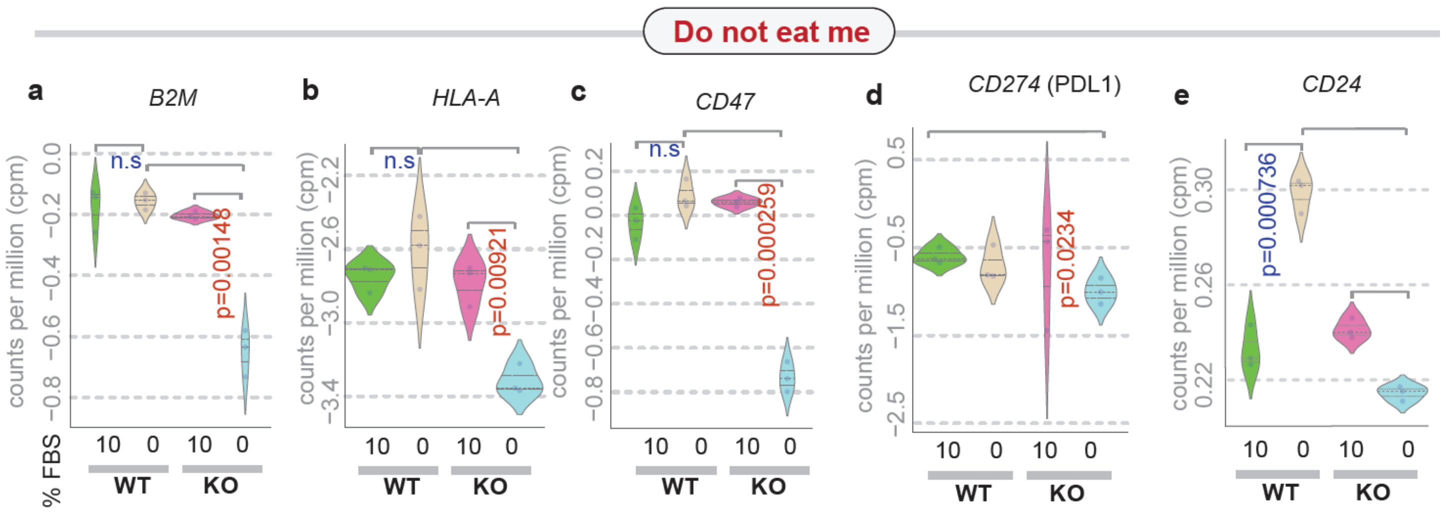


**Figure S6.** [Related to Figure 6]

**GIV-dependent growth signaling autonomy impacts the expression of immune checkpoint proteins.**

Violin plots show the levels of expression of various genes that encode immune checkpoint proteins in parental (WT) and GIV-depleted (GIV-KO) MDA MB-231 cells. See **Figure 6c-d** for the composite score for these genes.
